# Recapitulating memory B cell responses in a Lymphoid Organ-Chip to evaluate mRNA vaccine boosting strategies

**DOI:** 10.1101/2024.02.02.578553

**Authors:** Raphaël Jeger-Madiot, Delphine Planas, Isabelle Staropoli, Jérôme Kervevan, Héloïse Mary, Camilla Collina, Barbara F. Fonseca, Hippolyte Debarnot, Rémy Robinot, Stacy Gellenoncourt, Olivier Schwartz, Lorna Ewart, Michael Bscheider, Samy Gobaa, Lisa A. Chakrabarti

## Abstract

Predicting the immunogenicity of candidate vaccines in humans remains a challenge. To address this issue, we developed a Lymphoid Organ-Chip (LO chip) model based on a microfluidic chip seeded with human PBMC at high density within a 3D collagen matrix. Perfusion of the SARS-CoV-2 Spike protein mimicked a vaccine boost by inducing a massive amplification of Spike-specific memory B cells, plasmablast differentiation, and Spike-specific antibody secretion. Features of lymphoid tissue, including the formation of activated CD4+ T cell/B cell clusters and the emigration of matured plasmablasts, were recapitulated in the LO chip. Importantly, myeloid cells were competent at capturing and expressing mRNA vectored by lipid nanoparticles, enabling the assessment of responses to mRNA vaccines. Comparison of on-chip responses to Wuhan monovalent and Wuhan/Omicron bivalent mRNA vaccine boosts showed equivalent induction of Omicron neutralizing antibodies, pointing at immune imprinting as reported *in vivo*. The LO chip thus represents a versatile platform suited to the preclinical evaluation of vaccine boosting strategies.

## INTRODUCTION

Secondary lymphoid organs (SLO) and tertiary lymphoid structures (TLS) provide the adequate microenvironment for antigen (Ag)-specific lymphocytes to cooperate and develop efficient adaptive immune responses. Upon Ag stimulation, CD4+ T cell-independent B cells allow the rapid development of a first round of protective antibody responses against incoming pathogens. In a second stage, the collaboration between Ag-specific CD4+ T cells and Ag-specific B cells triggers the formation of transitory polarized areas called germinal centers (GC) where antibody responses are optimized through cycles of somatic hypermutation and class switch recombination (*1*). Upon Ag boosting, antibody-secreting cells matured in the GC leave SLO and peak in the circulation about one week later, before homing to effector tissues for terminal differentiation and sustained antibody production (*2*) .The matured antibodies, with increased affinity for Ag and broader functionalities, play a key role in limiting pathogen spread and provide long-term immunity against reinfection (*3*).

Deciphering human immune responses within SLO and TLS remains challenging. Animal models have provided most of the mechanistic knowledge on the maturation of Ag-specific immune responses, but they do not fully recapitulate human physiology and, hence, cannot accurately predict immunogenicity in human populations (*4*). The search for alternative preclinical systems has spurred the development of humanized mouse models, which have yielded valuable insights in terms of pathogenesis, but do not yet develop fully mature SLOs (*5*). Fine needle aspirations (FNA) from SLO of human volunteers represent a promising tool as they capture the native lymphoid environment (*6*)(*7*). However, FNA remain invasive and restrict the evaluation of immune functions due to the collection of a limited number of cells. In recent years, many efforts have focused on developing human cell-based models of SLO for studying adaptive immune responses *in vitro*. In a simple implementation, cocultures of human CD4+ T cells and B cells were used to demonstrate the key role of T follicular helper cells (Tfh), a specialized CD4+ T cell subset, in providing help for B cell maturation in the context of viral infections and autoimmune diseases (*8*, *9*). Research then aimed to recreate the three-dimensional (3D) environment typical of SLO, which promotes cell motility and intercellular interactions, thus increasing chances of encounter between Ag and Ag-specific cells (*10*). Explants of SLOs, consisting in blocks of adenoids or tonsils cultivated on collagen sponges, have been used successfully to evaluate HIV replication (*11*) and the early CD4+ T cell recall responses to pertussis toxin (*12*). However, tissue explants are not applicable to the study of antibody production due to their short half-lives. More recently, Wagar *and coll.* developed long-term cultures (≥2-3 weeks) of dissociated tonsil cells that spontaneously reassembled into lymphoid-like aggregates (*13*). These tonsil organoids demonstrated class switch recombination and antibody production upon stimulation with several viral vaccines, and thus proved useful to compare human immune responses to different vaccination modalities *in vitro* (*14*).

3D cultures based on peripheral blood mononuclear cells (PBMC) have the decisive advantage of allowing the assessment of human immune responses in large cohorts of vaccinated volunteers, without the need for invasive lymphoid tissue sampling. A 3D environment mimicking a lymphoid-like architecture can be provided by porous synthetic scaffolds or by a variety of hydrogels which are enriched in extracellular matrix (ECM) components such as collagen or fibrin (*15*, *16*). Researchers who focused on modeling the early steps of antigenic priming developed tissue-like 3D cultures where primed monocytes mature into dendritic cells (DC) upon Ag stimulation and emigrate through an endothelial barrier, before being collected and cocultured with purified T and/or B lymphocytes (*17*, *18*). This more physiological approach to DC priming was shown to recapitulate the effect of age on tuberculosis or influenza vaccination, with decreased T and/or B cell responses in cultures from elderly volunteers (*17*, *19*). Several groups have achieved an efficient differentiation of antibody-secreting B cells by bypassing the requirements for CD4+ T cell help through the inclusion of CD40L-expressing fibroblasts and cytokines in cultures (*20–22*). For instance, plasmablast differentiation was achieved in a 3D culture where B cells were cocultured with CD40L- and BAFF-expressing fibroblasts within a polyethylene-glycol hydrogel (*23*)(*20*). Functionalization of this hydrogel by integrin ligands markedly enhanced antibody secretion upon B cell receptor (BCR) ligation, emphasizing the importance of tissue-dependent signaling for plasmablast differentiation.

Microfluidic systems that ensure a continuous perfusion of 3D cultures further increase cell adhesion, migration, and differentiation (*24*), as physiological levels of shear stress promote integrin function and chemokine production (*25*, *26*). In addition, the continuous renewal of nutrients and controlled oxygenation levels in microfluidic devices enable cultures at high cellular densities that approach those observed in lymphoid organs (*16*)(*15*). Both the increased migration capacity and high cellular density facilitate the encounter of rare Ag-specific CD4+ T cells and B cells, emphasizing the potential of microfluidic systems for mimicking the GC environment. Using a microfluidic device where purified CD4+ T cells and B cells were cocultured within an ECM gel, Goyal *and coll.* showed that fluid perfusion could trigger the self-organization of these cells into lymphoid-like follicles (*28*). Further, a specific antibody response to an influenza vaccine could be detected in this system after autologous DC addition, without the need for extraneous CD40L or cytokine stimulation. Microfluidics technology thus opens the possibility of evaluating CD4+ T cell/B cell interactions in conditions that more closely mimic human physiology and that may better predict humoral responses *in vivo*.

The COVID-19 pandemic has emphasized the need for preclinical systems that enable a rapid evaluation of humoral responses elicited by candidate vaccines. Due to the successive emergence of SARS-CoV-2 variants and the implementation of various COVID vaccination strategies, the immunologic status of individuals is highly diverse and complex to decipher (*29–31*). The possibility to evaluate immune responses at a preclinical level in specific cohorts of vaccinated/infected individuals would clearly facilitate the development of optimal COVID booster vaccines. Hence, we developed a Lymphoid Organ-Chip (LO chip) model adapted to the testing of different COVID vaccine formulations. The LO chip relies on total PBMC embedded in a collagen-based matrix under slow perfusion within a microfluidic chip. This streamlined system promoted a potent amplification of Ag-specific memory B cells and neutralizing antibody production without the requirement for large blood samples nor purified immune cell populations. Importantly, the presence of myeloid cells allowed the capture and expression of mRNA vectored by lipid nanoparticles, enabling the assessment of responses to the new generation of COVID vaccines.

## RESULTS

### Amplification of Spike-specific B cell responses in the LO chip

To establish a 3D human lymphoid culture, we used a two-channel microfluidic chip developed by Emulate. The S1^®^ chip is composed of polydimethylsiloxane (PDMS), a polymer that enables gaseous exchanges, and is dynamically perfused by medium through connection to a microfluidic controller. The lower chip channel, devised as a tissue-like compartment, contains human PBMC seeded at a high concentration (6×10^8^/mL) within a collagen-based ECM, to mimic the high cellular densities achieved in lymphoid tissue. The upper channel is used as a vascular-like compartment to perfuse nutrients and Ag, which can reach the tissue-like compartment by diffusion through a porous membrane (Fig. 1A). The LO chip model was benchmarked using PBMC from healthy blood donors who had been previously exposed to SARS-CoV-2 and/or to COVID vaccines, based on their positive serology for the Spike Ag. To determine whether we could induce a Spike-specific recall B cell response for these donors, the LO chips were perfused for 6 days with a recombinant Spike protein derived from the ancestral SARS-CoV-2 Wuhan strain or with a control Ag, bovine serum albumin (BSA). At the end of the incubation period, the ECM was recovered and digested to allow the analysis of cell phenotype by flow cytometry (see gating strategy in Fig. S1).

**Figure 1.**
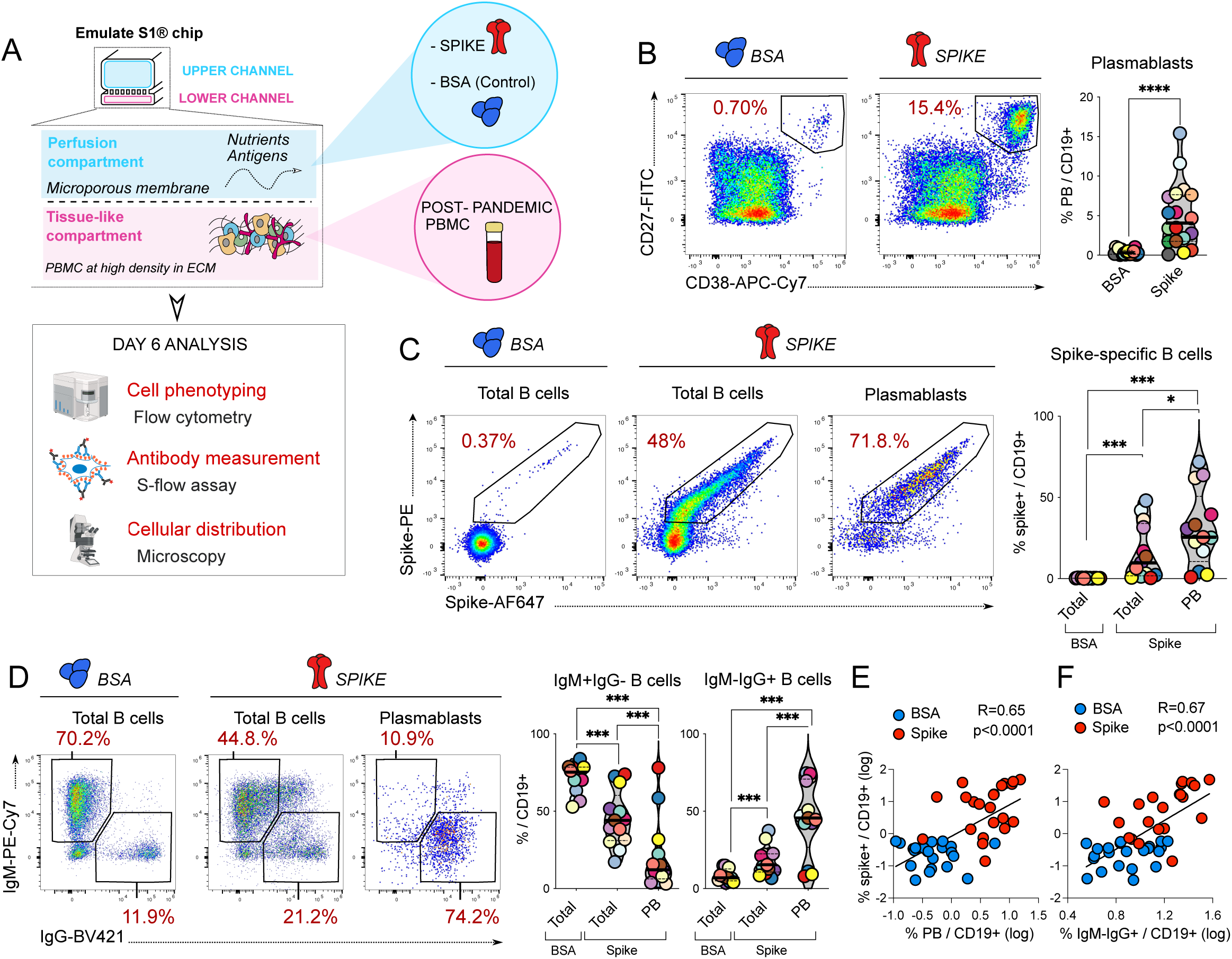
Amplification of Spike-specific memory B cells in the LO chip. (A) Experimental design: The LO chip consists in an S1^®^ microfluidic (Emulate), where the top channel is perfused with antigen (Spike protein or BSA control), while the bottom channel is seeded with PBMC at high-density in a collagen-based extracellular matrix. PBMC were obtained from post-pandemic donors positive for SARS-CoV-2 Spike antibodies. (B) Representative dot plots of CD27 and CD38 staining in CD19+ B cells (left) and frequency of CD27hiCD38hi plasmablasts (PB) among B cells recovered from LO chips at day 6. (C) Representative analysis of Spike-specific B cells labeled with two different fluorescent Spike proteins (left) and frequency of Spike-specific cells among total B cells and plasmablasts (right). (D) Representative analysis of IgM and IgG expression in total B cells and PB (left) and frequency of IgM+IgG- and IgM-IgG+ subsets among total B cells and PB (right). (E-F) Positive correlation between the frequency of Spike-specific B cells (y axis) and the frequency of PB (E) or of IgM-IgG+ B cells (F). The linear correlation coefficient R and the associated P value are reported. (B-D) Each color represents an independent donor. Differences were evaluated with a Wilcoxon matched pairs test. *p <0.05; **p < 0.01, ***p < 0.001, ****p < 0.0001.

Upon Spike perfusion, the frequency of CD27hiCD38hi plasmablasts (PB) increased among CD19+ B cells for 16 out of 18 donors tested (Fig. 1B left), with a median PB percentage rising from 0.3% in the BSA control to 4.1% in the Spike condition (Fig. 1B right). We then asked whether these signs of B cell maturation reflected an Ag-specific B cell response. Spike-specific B cells were detected by labeling with a fluorescently labeled Wuhan Spike protein, using the same protein labeled with two distinct fluorochromes to ensure low background staining (Fig. 1C left). A marked increase in Spike-specific B cells was observed in LO chips from 12 out of 13 donors tested, with a median specific B cell percentage rising from 0.3% to 9.8% of total B cells upon Spike stimulation (Fig.1 C right). This corresponded to a median 32.7-fold increase, highlighting the massive amplification of Spike-specific B cells after a 6-day culture in the LO chip. The frequency of Spike-specific B cells was further enriched in the PB population compared to the total B cell population, reaching a median percentage of 25.5% (Fig. 1C right), and suggesting that a high fraction of antibody-secreting B cells were Ag-specific.

As the GC reaction is known to trigger immunoglobulin class switch recombination (*1*), we measured the frequency of B cells expressing the IgM and IgG immunoglobulin isotypes (Fig. 1D left). We observed a shift from a predominantly IgM+IgG-B cell population to a mixed population with an increased frequency of IgM-IgG+ B cells, from a median of 6.9% in the BSA condition to 15.4% upon Spike stimulation. The acquisition of an IgM-IgG+ phenotype was more marked in PB, with a median of 45.6% cells expressing surface IgG in this population after Spike treatment (Fig. 1D right). Thus, antigenic stimulation in the LO chip promoted IgG expression in a significant fraction of B cells. The frequency of Spike-specific B cells showed a strong correlation with that of PB (Fig. 1E; R=0.65, P<0.0001) and with that of IgM-IgG+ B cells (Fig. 1F; R=0.67, P<0.0001), suggesting that a coordinated maturation of the B cell response to the Spike antigen took place in the LO chip.

### Production of Spike-specific neutralizing antibodies in the LO chip

To determine whether Spike-specific B cell maturation resulted in detectable antibody production, we measured Spike-binding IgG antibodies using the S-flow assay (*32*). This cell-based assay, which measures the amount of human IgG bound at the surface of Spike-expressing 293-T cells (Fig. 2A), has the advantage of detecting antibodies specific for the Spike in its native conformation and of being more sensitive than a classic ELISA. The production of Spike-specific IgG was induced for 12 out of 14 donors in Spike-stimulated compared with BSA-stimulated LO chips, with an increase in median MFI (mean fluorescent intensity) from 1.7 x 10^3^ to 60.3 x 10^3^ (Fig. 2B; P<0.001). These antibody measurements were done in the solution recovered from the chip after non-enzymatic ECM digestion, as the antibody concentrations measured in the chip effluent were low (Fig. S2A), suggesting that secreted antibodies were retained in the ECM. As expected, the amount of Spike-binding IgG recovered from the ECM showed a strong correlation with the frequency of Spike-specific B cells (Fig. 2C; R=0.77, P=0.0002). Spike-specific IgA production was also induced upon Spike stimulation 7 of 14 donors tested (Fig. S2B; P<0.01), reinforcing the notion that class switch recombination took place in the LO chip.

**Figure 2.**
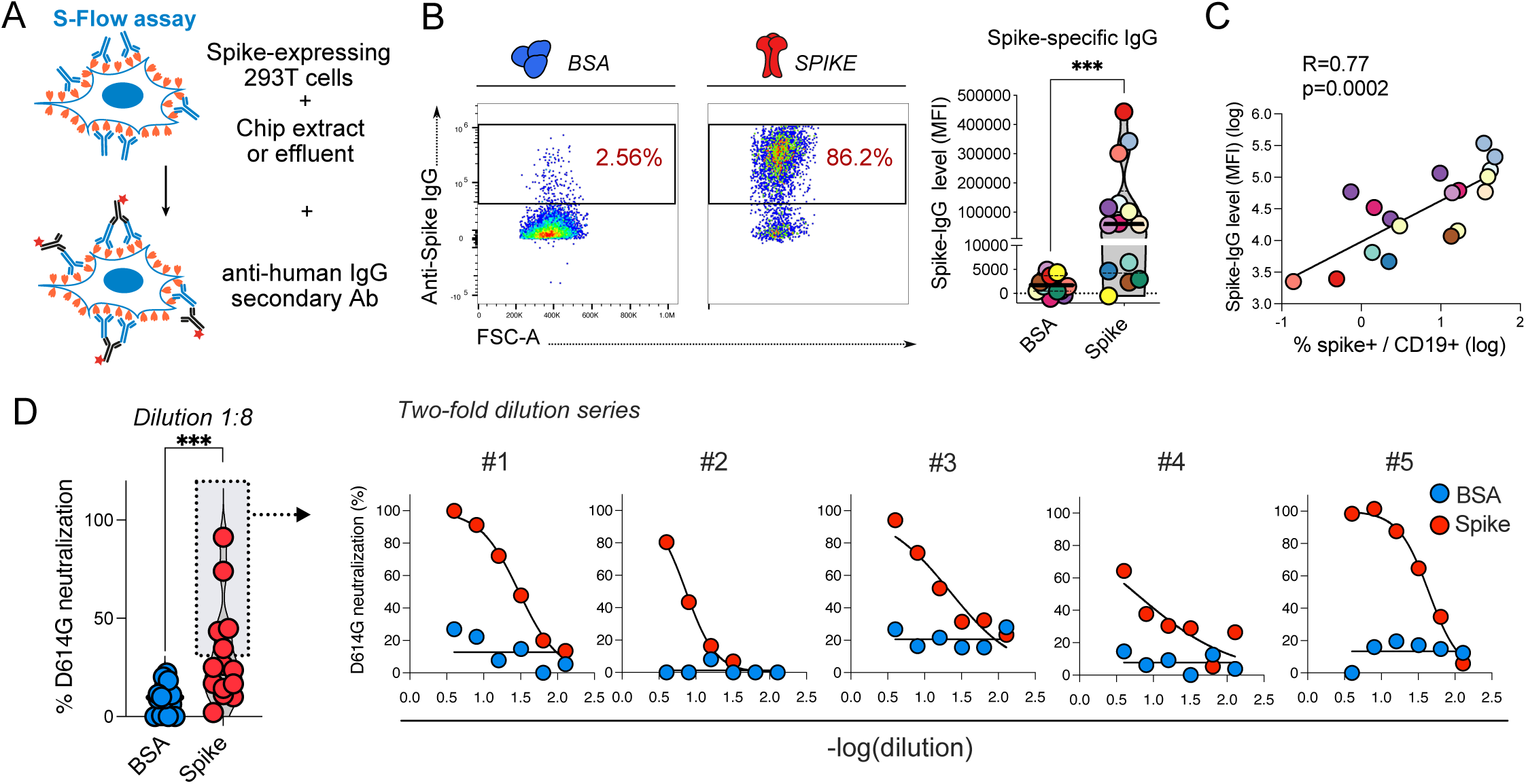
Early production of neutralizing antibodies in the LO chip. (A) The production of Spike-binding antibodies was assessed using the S-flow assay, where Spike-expressing 293T cells are exposed to chip extracts, and the binding of Spike-specific antibodies is revealed with a secondary anti-human IgG antibody. (B) Representative staining of control or Spike-specific IgG bound to Spike-expressing 293T cells (left) and mean fluorescent intensity (MFI) of chip-derived IgG bound to Spike-expressing 293T cells (right). (C) Positive correlation between Spike-specific IgG MFI and the frequency of Spike-specific B cells. The linear regression correlation coefficient R and the associated P-value are reported. (B, C) Each color represents an independent donor. (D) The neutralizing capacity of antibodies recovered from LO chips at day 6 was assessed for n=13 donors by the S-Fuse assay (left). The dose response of D614G neutralization in function of chip extract dilution is reported for the 5 donors who had a positive neutralization response at the 1/8 dilution (right). (B, D) Differences were evaluated with a Wilcoxon matched pairs test.; *p <0.05; **p < 0.01, ***p < 0.001, ****p < 0.0001.

We next evaluated the neutralizing activity of the antibodies produced in the LO chip upon Spike stimulation. To this goal, we used the S-Fuse assay, which consists in measuring the inhibition of infection in U2OS-ACE2 cells inoculated by SARS-CoV-2, using a GFP-split cell system (*33*). Neutralization activity was determined by the capacity of antibodies to limit infection and syncytia formation, as measured by a decrease in the GFP signal. Neutralizing activity against the SARS-CoV-2 strain D614G (a variant very close to the ancestral strain) could be detected or 5 out of 13 donors tested in the ECM solution diluted at 1:8 (Fig. 2D, left). Analyses of positive ECM samples showed a dose-dependent response upon serial dilutions (Fig. 2D, right), with a measured ID_50_ (dilution inhibiting 50% of infections) in the 6-44 range. Thus, neutralizing antibody production could be achieved in the LO chip even after a short culture duration of 6 days.

### Preferential egress of Spike-specific PB from the LO chip

In the context of a recall response, memory B cells can either differentiate rapidly into antibody secreting PB and plasma cells or adopt an activated memory B cell phenotype able to enter the GC reaction for a further round of B cell response maturation. B cells that differentiate directly into PB/plasma cells tend to have a higher BCR affinity for Ag compared to those that retain a memory phenotype (*1*). To determine whether these features were recapitulated in the LO chip, we first analyzed the phenotype of CD19+ B cells in more detail by defining 5 subsets based on CD27 and CD38 expression, as described (*34*). A classic distribution of B cell subsets was observed in LO chips, with the presence of double negative (DN) B cells (CD27-CD38-), memory B cells (CD27medCD38-), activated memory B cells (CD27medCD38med), unswitched B cells (CD27-CD38med), and PB (CD27hiCD38hi), as shown in Fig. 3A. Analysis of Spike-stimulated chips showed a significant increase in both PB and activated memory B cells in the Spike-specific population compared to the non-specific population from the same chip (Fig. 3B). In contrast, the resting memory B cell subset and the unswitched B cell subset (which preferentially express IgM; see Fig. 3C below) declined in the Spike-specific population, while the DN subset remained unchanged. Thus, Spike-specific B cells transitioned to more mature states and could adopt either an activated memory or a PB cell fate, like the differentiation pathways documented in lymphoid organs.

**Figure 3:**
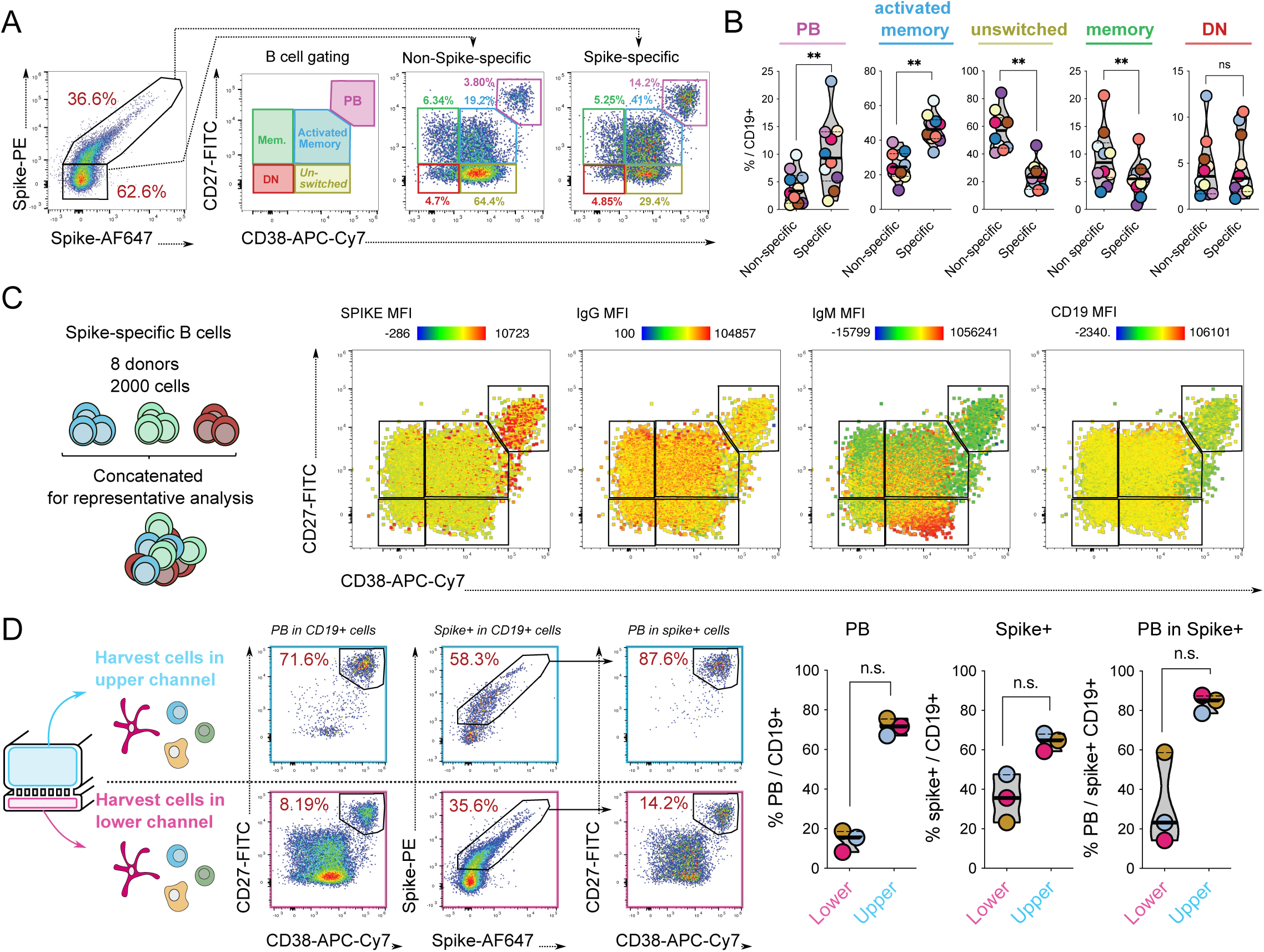
Differentiation of high-affinity plasmablasts in the LO chip. (A) CD19+ B cells were subdivided in several subsets: CD27hiCD38hi plasmablasts (PB), CD27medCD38med activated memory B cells, CD27-CD38med unswitched B cells, CD27medCD38-memory B, and CD27-CD38-double negative (DN) B cells. A representative example of subset frequencies in non-Spike-specific and Spike-specific B cells is shown. (B) Comparison of B cell subset frequencies in non-Spike-specific and Spike-specific B cells. (C) Flow cytometry files corresponding to Spike-specific B cells from 8 donors were concatenated (left) and the mean fluorescent intensities (MFI) corresponding to the binding level of the Spike and the expression of IgG, IgM, and CD19 were mapped by pseudocolor on the different B cell subsets defined by the CD27/CD38 parameters (right). (D) Cell harvesting scheme (left) and representative examples of B cell phenotyping on cell recovered from the upper (top line) and lower (bottom line) LO chip compartments (middle plots). Frequencies of PB in CD19+ B cells, Spike-specific B cells in CD19+ cells, and PB in the Spike-specific B cell subset (right) are reported for the upper perfusion compartment (Upper) and the lower tissue-like compartment (Lower). (B, D) Each color represents an independent donor. Differences were evaluated with a Wilcoxon matched pairs test; *p <0.05; **p < 0.01; n.s. not significant.

We then mapped the intensity of Spike-PE binding onto the different B cell subsets within the Spike-specific population (Fig. 3C, right plot). For this analysis, an identical number of Spike-specific B cells obtained from 8 donors were concatenated (Fig. 3C, left), resulting in a representative Spike-binding pattern. Interestingly, the PB subset showed a higher intensity of Spike binding as compared to the 4 other B cell subsets, which suggested a higher avidity of PB for the Spike protein. In contrast, expression of the BCR (both IgG and IgM) and of the coreceptor CD19 were lower in PB compared to the other 4 subsets (Fig. 3C, middle and right plots), consistent with a transition towards an antibody-secreting phenotype. As the higher Spike-binding capacity of PB could not be ascribed to increased BCR expression, these observations suggested that the PB that differentiated in the LO chip had a higher affinity for the Spike antigen compared to other B cell subsets.

Another feature of SLO is that differentiated PB exit the lymphoid tissue to join the lymph and blood circulation before homing to their supportive niches in the bone marrow or peripheral sites, while activated memory B cells are retained in GC (*3*). We noted that a fraction of cells seeded into the lower channel of the LO chip egressed to the upper channel during culture after Spike stimulation. To determine whether this phenomenon reflected PB exit from the tissue-like compartment, we recovered cells separately from the upper and lower channels at the end of the 6-day Spike-stimulation (Fig. 3D, left schematic). Phenotyping showed that B cells that egressed were highly enriched in PB (median= 71.6% PB in upper channel) and were also enriched in Spike-specific cells (median=64.8% Spike+ B cells in upper channel; Fig. 3D, left and middle graphs). Consistently, most of the egressed Spike-specific B cells were PB (median=85.1% PB in Spike+ B cells from upper channel; Fig. 3D right graph). Taken together, these findings indicate that B cell maturation in the LO chip recapitulates important features observed in SLO, with an increased affinity for antigen in PB compared to memory B cells, and a preferential egress of PB from lymphoid tissue.

### Amplification of Ag-specific CD4+ T cells and formation of CD4+ T cell/B cell clusters in the LO chip

CD4+ T cell help plays a key role in the maturation of the B cell response. We asked whether we could detect activated CD4+ T cells with B cell helper capacity, based on the co-expression of the activation marker CD38 and the costimulatory receptor ICOS (*35*, *36*). A marked increase in CD38+ICOS+ CD4+ T cells was observed in cells recovered from Spike-stimulated LO chips at day 6, compared to BSA-stimulated chips (Fig. 4A). The frequency of CD38+ICOS+ CD4+ T cells correlated with that of PB, suggesting a concomitant induction of CD4+ T cell activation and B cell maturation (Fig. 4B; R=0.70, P<0.0001). We restimulated cells recovered from chips with a pool of overlapping peptides spanning the Spike antigen to evaluate the specific CD4+ T cell response. A significant induction of CD4+ T cells producing TNF-α, IFN-γ, or both cytokines was observed in the Spike-stimulated sample (Fig. 4C). IL-2 production was also induced in CD4+ T cells upon Spike stimulation (Fig. 4D) and was enriched in the subset of cells that also produced IFN-γ and TNF-α (Fig. 4E), showing the induction of polyfunctional CD4+ T cells. Thus, specific CD4+ T cells were amplified in parallel to specific B cells in Spike-stimulated LO chips.

**Figure 4.**
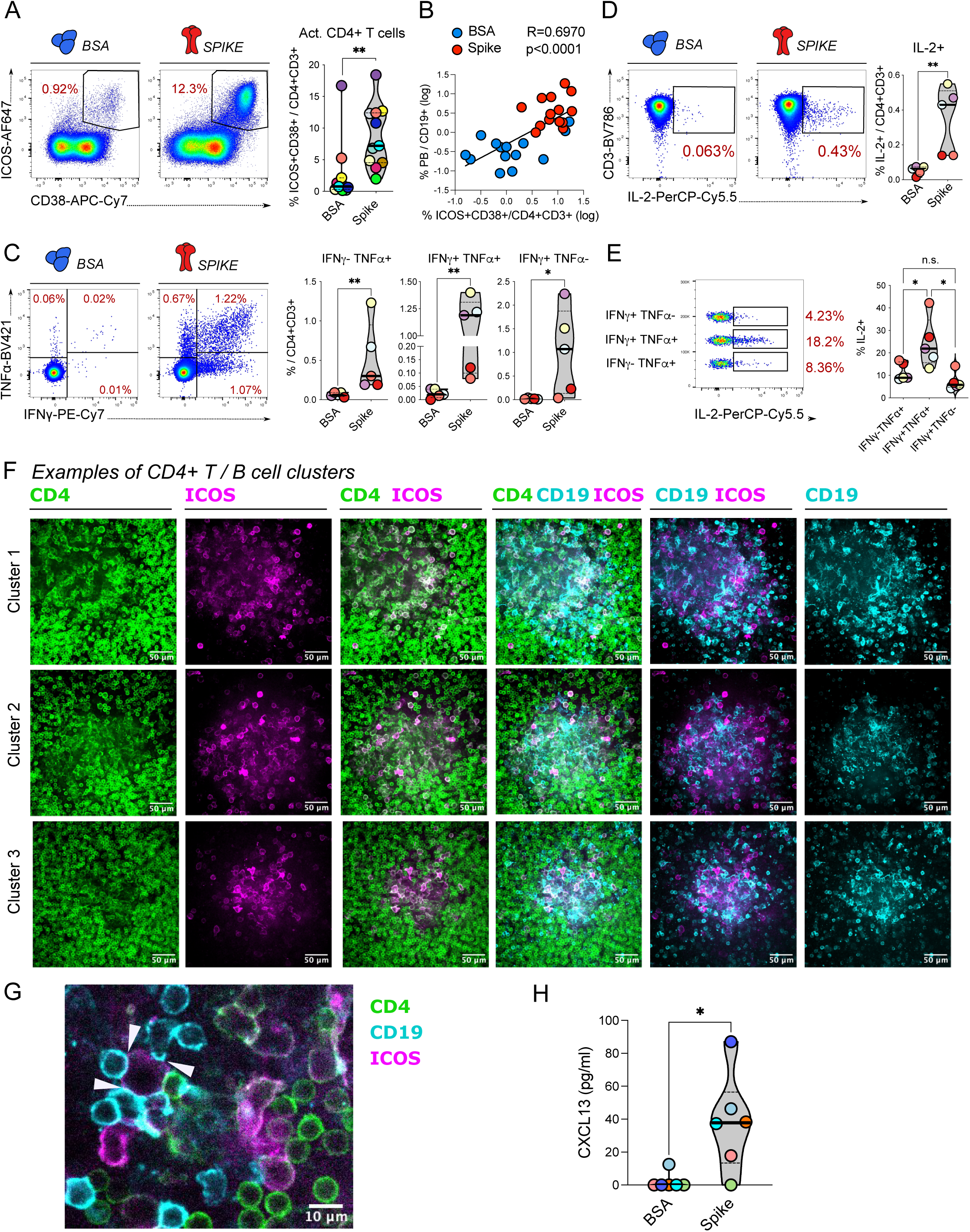
Amplification of Spike-specific CD4+ T cells and spatial association of costimulatory CD4+ T cells with B cells in the LO chip. (A) Representative analysis of ICOS and CD38 expression in CD3+CD4+ cells at day 6 after antigenic stimulation (left) and frequency of CD38+ICOS+ activated cells among CD3+CD4+ T cells (right). (B) Positive correlation between the frequency of CD38+ICOS+ cells among CD4+ T cells and the frequency of CD27hiCD38hi plasmablasts (PB) among B cells. The linear correlation coefficient R and the associated P value are reported. (C) Representative analysis of TNF-α and IFN-γ expression in CD4+ T cells after overnight restimulation with a Spike-peptide pool following 6 days of LO Chip culture (left), and frequency of Spike-specific cells expressing TNF-α, IFN-γ, or both among CD4+ T cells (right). (D) Representative analysis of IL-2 expression in CD4+ T cells after overnight restimulation with a Spike-peptide pool following 6 days of LO Chip culture (left) and frequency of Spike-specific cells expressing IL-2 among CD4+ T cells (right). (E) Frequency of IL-2 expression in Spike-specific CD4+ T cell subsets defined by TNF-α and IFN-γ expression. (F) Confocal microscopy image of a 6-day LO chip culture perfused with the Spike antigen. B cells were stained with CD19 (cyan), CD4+ T cells with CD4 (green), and activated CD4+ T cells with ICOS (magenta). A representative image of a CD4+T cell/B cell cluster is shown at x40 magnification, using a Z stack projection on maximal intensity, with single and merged channels. (G) Focus on an ICOS+ CD4+ T cell showing synapses (arrowheads) with multiple B cells. (H) ELISA measurement of CXCL13 production in LO chip extracts obtained at day 6. (A, C-E, H) Each color represents an independent donor. Differences were evaluated with a Mann-Whitney test; *p < 0.05; **p < 0.01; n.s. not significant.

To investigate whether CD4+ T cells and B cells spatially interact in the LO chip, we analyzed the chips by fluorescence microscopy at day 6. Low magnification imaging of the lower channel revealed that Spike stimulation induced the formation of large cell clusters where CD19+ B cells colocalized with CD4+ T cells, while cells remained more dispersed upon BSA stimulation (Fig. S3 A-B). B cells aggregated at high density within the clusters, while CD4+ T cell density appeared more variable. Interestingly, CD4+ T cells expressing the costimulatory marker ICOS were highly enriched within the clusters (Fig. S3A), suggesting the formation of GC-like areas where activated CD4+ T cells interacted with B cells. This notion was confirmed by the observation that the proliferation marker Ki67 was confined to the clusters (Fig. S3B), suggesting that CD4+ T cell/B cell interactions promoted cellular proliferation. Analysis of the clusters at higher magnification confirmed the enrichment of CD4+ T cells expressing ICOS within these areas (Fig. 4F). The ICOS+ CD4+ T cells had an ameboid shape and were enlarged compared to their ICOS-counterparts, consistent with cellular activation in the ICOS+ population. The ICOS+ CD4+ T cells could form tight contacts with one or more B cells, suggestive of immunological synapse formation (Fig. 4G). Further, Spike stimulation induced the secretion of the CXCL13 chemokine, a marker of GC activity (Fig. 4H) (*37*). Thus, specific antigen stimulation in the LO chip promoted the formation of cellular clusters that recapitulated several features of GC, including enrichment in CD4+ T cell/B cell interactions, immune activation, cellular proliferation, and CXCL13 secretion.

### Dynamic culture conditions in the LO chip promote Spike-specific responses

To investigate the effect of continuous fluid perfusion on antigen-specific responses, we compared dynamic cultures in the LO chip to static 3D cultures. To this goal, we seeded an equal number of PBMC (10^7^) within an equal amount of ECM gel (17 µL) either in the lower channel of an LO chip or as a drop in a well of a classic tissue culture plate (Fig. 5A). After stimulation with Ag, we could compare the reactivity of PBMC grown at a similar density in 3D, either in static or in dynamic conditions. Of note, we chose not to perform the static culture within a microfluidic chip, as we reasoned that the restricted diffusion of nutritive medium through the channels would be insufficient to sustain a high-density culture. With our setup, Spike stimulation resulted in a limited induction of PB at day 6 for the in-gel culture, while PB induction was significantly higher for the in-chip culture (Fig. 5B; P<0.05). We also observed that simple 2D cultures without ECM resulted in even lower PB induction than in-gel cultures, consistent with a beneficial effect of extracellular matrix components and/or of a 3D spatial organization (Fig. S4).

**Figure 5.**
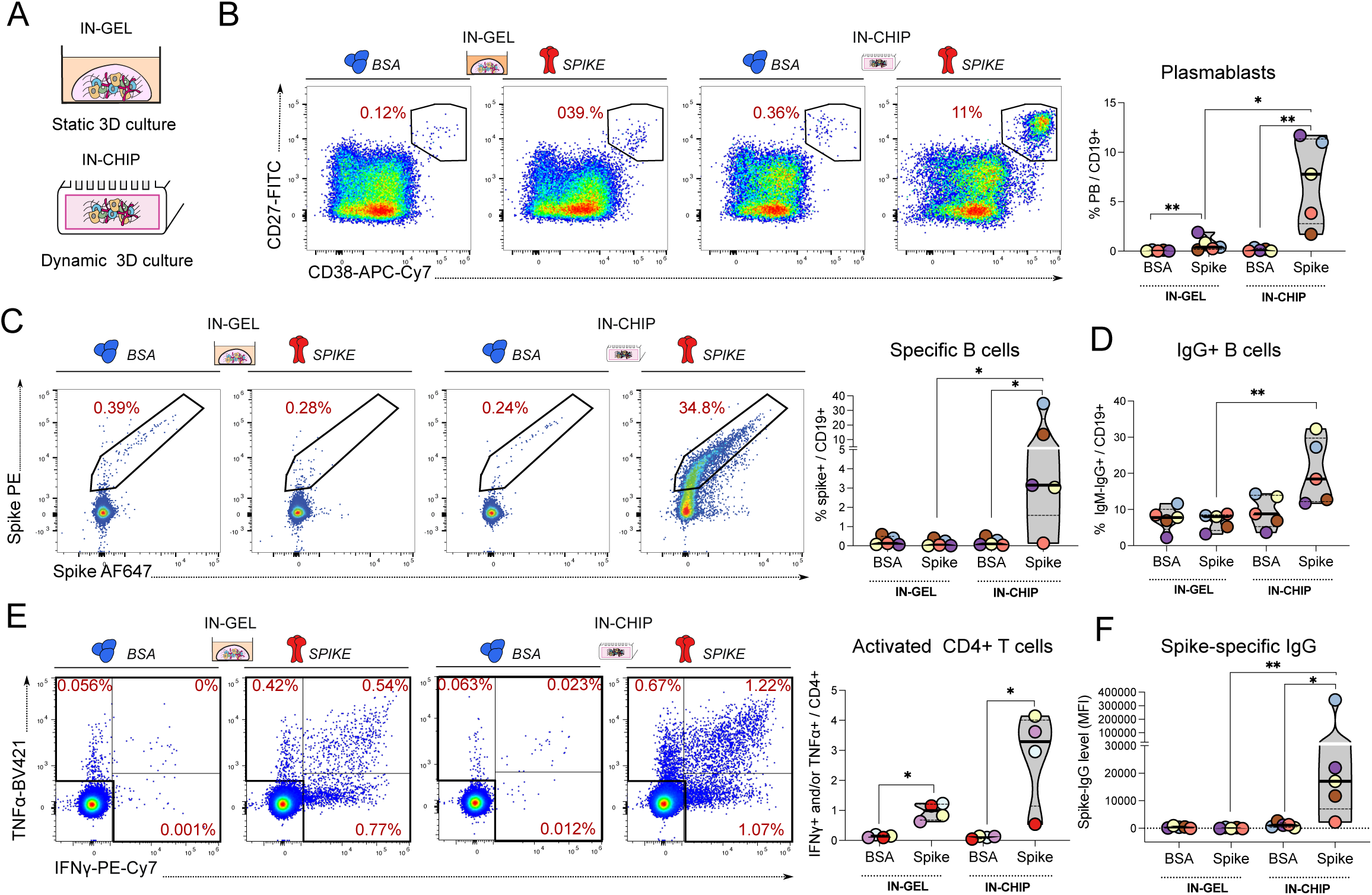
Dynamic fluid flow increases Spike-specific CD4+ T cell and B cell responses in the LO chip. (A) Experimental setup: the same number of PBMC were cultivated at the same cellular density for 6 days either in the perfused LO chip system or in a gel-droplet placed in a 24-well plate, using the same concentration of BSA and Spike antigens. (B) Representative analysis of CD27 and CD28 expression in B cells (left) and frequency of CD27hiCD38hi plasmablasts (PB) among B cells recovered from gel droplets (in-gel) or from LO chips (in-chip). (C) Representative analysis of fluorescent Spike labeling of B cells (left) and frequency of Spike-specific cells among CD19+ B cells (right) obtained in the gel-droplet culture and the LO chip. (D) Frequency of IgM-IgG+ cells among CD19+ B cells obtained in the gel-droplet culture and the LO chip. (E) Representative analysis of TNF-α and IFN-γ expression in CD4+ T cells after overnight restimulation with a Spike-peptide pool after 6 days of culture (left), and frequency of Spike-specific cells expressing TNF-α and/or IFN-γ (single and double positive cells) among CD4+ T cells (right) obtained in the gel-droplet culture and the LO chip. (F) Mean fluorescent intensity (MFI) of Spike-specific IgG bound to Spike-expressing 293T cells (S-flow assay). (B-F) Each color represents an independent donor. Differences were evaluated with a Mann-Whitney test; *p < 0.05; **p < 0.01.

We then evaluated the effect of fluid perfusion on Spike-specific responses. The amplification of Spike-specific B cells remained undetectable in-gel, while it was detected in 4 out of 5 donors tested in-chip (Fig. 5C). Similarly, an increase in the percentage of IgM-IgG+ B cells upon Spike stimulation was detected in-chip, but not in-gel (Fig. 5D; P<0.01). Analyses of Spike-specific CD4+ T cell responses upon peptide restimulation showed that specific CD4+ T cells producing TNF-α and/or IFN-γ could be significantly induced both in-gel and in-chip, though the induction reached higher levels in-chip (Fig.5E). Thus, B cells appeared to have a more stringent requirement than CD4+ T cells for dynamic culture conditions. This notion was further confirmed by the analysis of Spike-binding IgG production, as only B cells from dynamic LO chip cultures produced Spike-specific antibodies at day 6, while B cells from static 3D cultures failed to do so Fig. 5F). These findings show that microfluidic perfusion in the LO chip significantly contributes to the amplification of B and CD4+ T cell memory responses.

### Myeloid antigen presenting cells (APC) are preferentially targeted by mRNA vaccines in the LO chip

We then explored the potential of the LO chip to evaluate recall responses to mRNA vaccines, as this new vaccination modality has proved instrumental in the rapid deployment and success of COVID vaccines (*38*). Studies in mouse and non-human primate models suggest that mRNA vaccines vectored by lipid nanoparticles (LNP) are primarily expressed by APC *in vivo*, and that mRNA expression peaks at early time points (*39–41*). Therefore, we perfused LO chips with the Spike-encoding mRNA vaccine mRNA-1273 (Moderna), harvested the chips at an early time point (day 2), and analyzed the status of different APC populations including CD11c+ HLA-DRhi DC, CD14+ monocytes, and CD19+ B cells (see gating strategy in Fig. S5). BSA protein stimulation was kept as a reference in these experiments.

Spike expression was detected using the human IgG1 mAb s102 directly coupled to the AF647 fluorophore (*33*, *42*), with the control consisting in the unrelated human IgG1 mAb GO53 coupled to the same fluorophore (Fig.6A, top and middle rows). Interestingly, mRNA-1273 treatment caused a significant increase in the frequency of Spike+ cells among DC and monocytes, but not among B cells nor CD4+ T cells (Fig. 6A, bottom row), suggesting that only myeloid cells were efficiently targeted by the mRNA vaccine. We also noted a limited but significant increase of DC and monocytes labeled with the control GO53 antibody after mRNA-1273 treatment, suggesting that the mRNA-LNP was sufficient to induce an activation of APC that increased non-specific antibody binding, consistent with the reported self-adjuvanting effect of this type of vaccine (*43*). To verify the nature of cells capable of expressing mRNA-LNP in the LO chip, we used mRNA-LNP that coded either for the green fluorescent protein (GFP) or for the control protein ovalbumin (OVA) and that had similar lipid composition and mRNA modifications as those of Moderna mRNA vaccines (Fig. 6B and S6). After 2 days of GFP mRNA-LNP perfusion, expression of GFP could be detected in DC and monocytes, but not in B cells nor CD4+ T cells, consistent with findings obtained with the mRNA-1273 vaccine.

**Figure 6.**
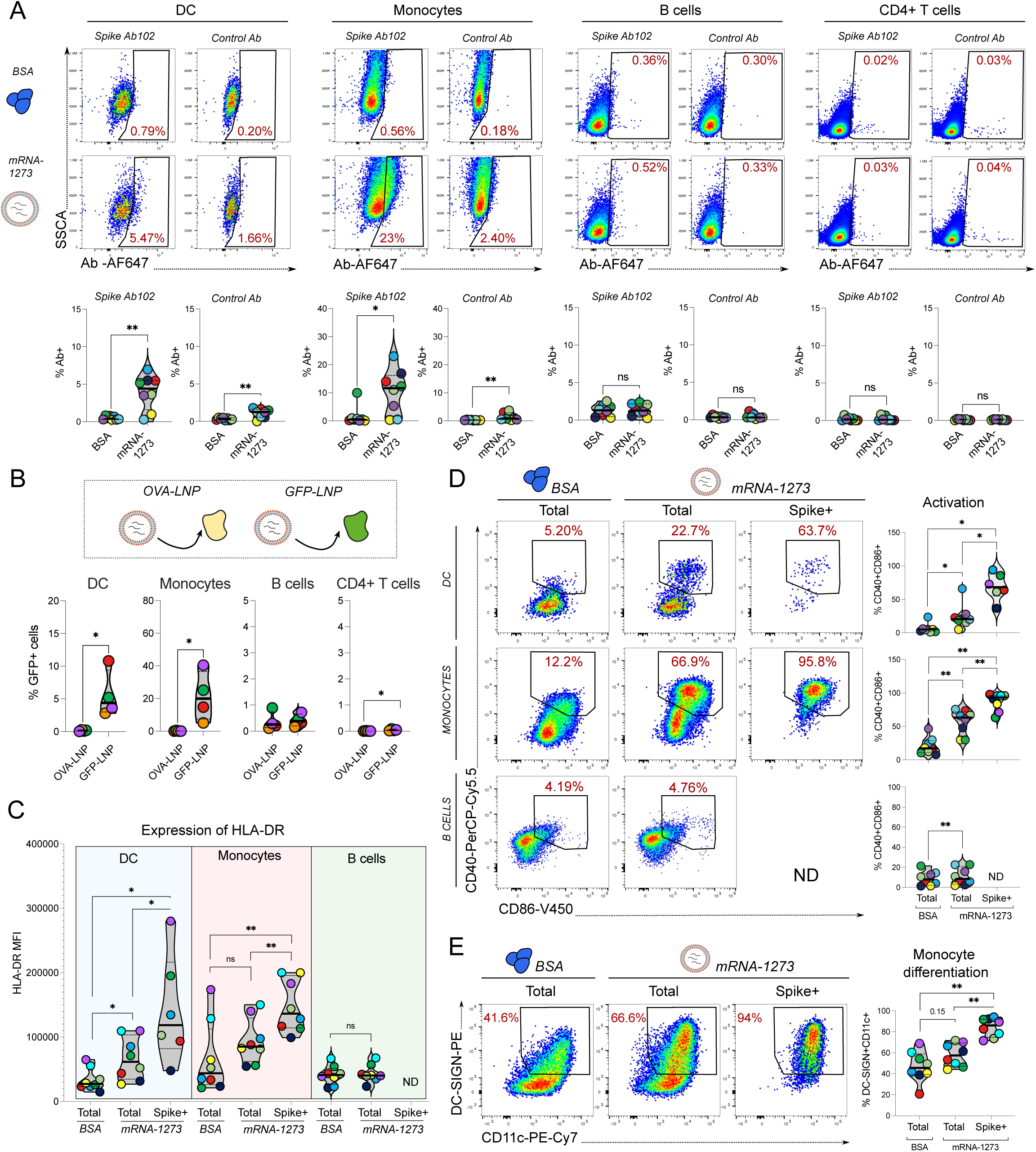
Preferential Spike expression and activation in myeloid cells upon mRNA vaccine stimulation in the LO chip. (A) After 2 days of perfusion with BSA (top row) or the mRNA-1273 vaccine (middle row), cells were harvested from the LO chip and stained with a panel focusing on myeloid cells. Spike expression was assessed with a directly labeled anti-Spike monoclonal antibody (Ab102-AF647; left panels for each cell type) and controlled using an irrelevant isotype-matched IgG1Ab (mGO53-AF647; right panels) Representative analyses at day 2 of Spike expression (top and middle rows) and frequency of Spike+ cells among dendritic cells (DC), monocytes, B cells and CD4+ T cells (bottom row) are reported. (B) LO chips were perfused for 2 days with lipid nanoparticles (LNP) containing mRNAs coding either for a control protein (ovalbumin, OVA) or for the green fluorescent protein (GFP) reporter. The frequency of GFP+ cells among DC, monocytes, B cells, and CD4+ T cells is reported. Differences were evaluated with a Mann Whitney test; *p <0.05. (C) Mean fluorescent intensity (MFI) of HLA-DR expression in total DC, monocytes, and B cells and the Spike-expressing subset (Spike+) of these cell types, after 2 days of stimulation with either BSA or the mRNA-1273 vaccine. (D) Representative analysis of CD40 and CD86 expression among DC, monocytes, B cells and their Spike-expressing subsets (left) and frequency of double positive CD40+CD86+ activated cells among these cell populations (right). ND: not detectable. (E) Representative analysis of CD11c and DC-SIGN expression among total monocytes and the Spike+ monocyte subset (left) and frequency of CD11c+DC-SIGN+ differentiated cells among total monocytes and the Spike+ monocyte subset (right). (A-E) Each color represents an independent donor. (A, C, D, E) Differences were evaluated with a Wilcoxon matched pairs test; *p <0.05; **p < 0.01; ns: not significant.

The analysis of activation markers at day 2 after mRNA-1273 perfusion showed a significant increase in HLA-DR expression in DC and a trend for increase in monocytes, while no increase was detected in B cells (Fig. 6C). The induction of HLA-DR was more marked in DC and monocytes that expressed the Spike (populations labeled as Spike+ in Fig. 6A), suggesting that these cells could be directly activated by mRNA LNP capture and/or expression. The expression of the costimulation markers CD40 and CD86 was also strongly induced in DC and monocytes treated with mRNA-1273, while it was induced to a lower extent in B cells (Fig. 6D). Further analysis showed a marked enrichment of CD40/CD86 coexpression in the subset of Spike+ DC and monocytes, confirming that myeloid cells which captured and expressed the mRNA-1273 LNP had the most prominent signs of activation. In addition, Spike+ monocytes showed an induction of typical DC-markers, namely CD11c and DC-SIGN (Fig. 6E), suggesting that mRNA vaccine capture could trigger monocyte differentiation towards a DC phenotype. Taken together, these findings indicated that the DC and monocytes present in the LO chip were competent at capturing and expressing mRNA-LNP vaccines. Further, these myeloid cells upregulated activation and differentiation markers, suggesting the induction of antigen presentation capacity.

### Efficient B cell responses to an mRNA vaccine boost in the LO chip

We then analyzed the induction of Spike-specific B cell recall responses upon mRNA vaccine stimulation in LO chips harvested at day 6 (Fig. 7A). A significant induction of PB was observed upon mRNA-1273 stimulation (Fig. 7B, P<0.01), though PB frequency appeared lower than that observed with Spike protein stimulation (difference not significant). Similarly, Spike-specific B cells were induced by mRNA-1273 stimulation (Fig. 7C, P<0.01), with a non-significant trend for lower specific B cell frequencies compared to samples stimulated with the Spike protein. Interestingly, we observed that the phenotype of amplified specific B cells differed between the two stimulation conditions, with a predominance of PB among the specific B cells induced by the mRNA vaccine (Fig. 7D; median PB in Spike+ cells: 41.8% for mRNA-1273 vs 12.8% for Spike, P<0.05). Note that only samples with a sufficiently high frequency of Spike+ cells were kept for this analysis (4 out of 7 samples in each group, with ≥ 120 Spike+ cells), to ensure a robust phenotypic analysis. In line with PB frequency, the production of Spike-specific IgG was higher upon the perfusion of the mRNA-based vaccine than of the Spike protein (Fig.7E; P<0.05). The production of Spike-specific IgA induced by the mRNA vaccine was also significantly higher (Fig. 7F, P<0.05), suggesting the induction of class switch recombination. The finding of increased antibody production appeared generalizable to another mRNA vaccine, as shown by a trend for higher Spike-specific IgG production upon stimulation by the BNT162b2 vaccine (Pfizer-BioNTech) compared to the Spike protein (Fig. 7G). Taken together, these findings indicated that mRNA vaccine boosting in the LO chip resulted in the induction of an efficient B cell recall response dominated by mature PB.

**Figure 7.**
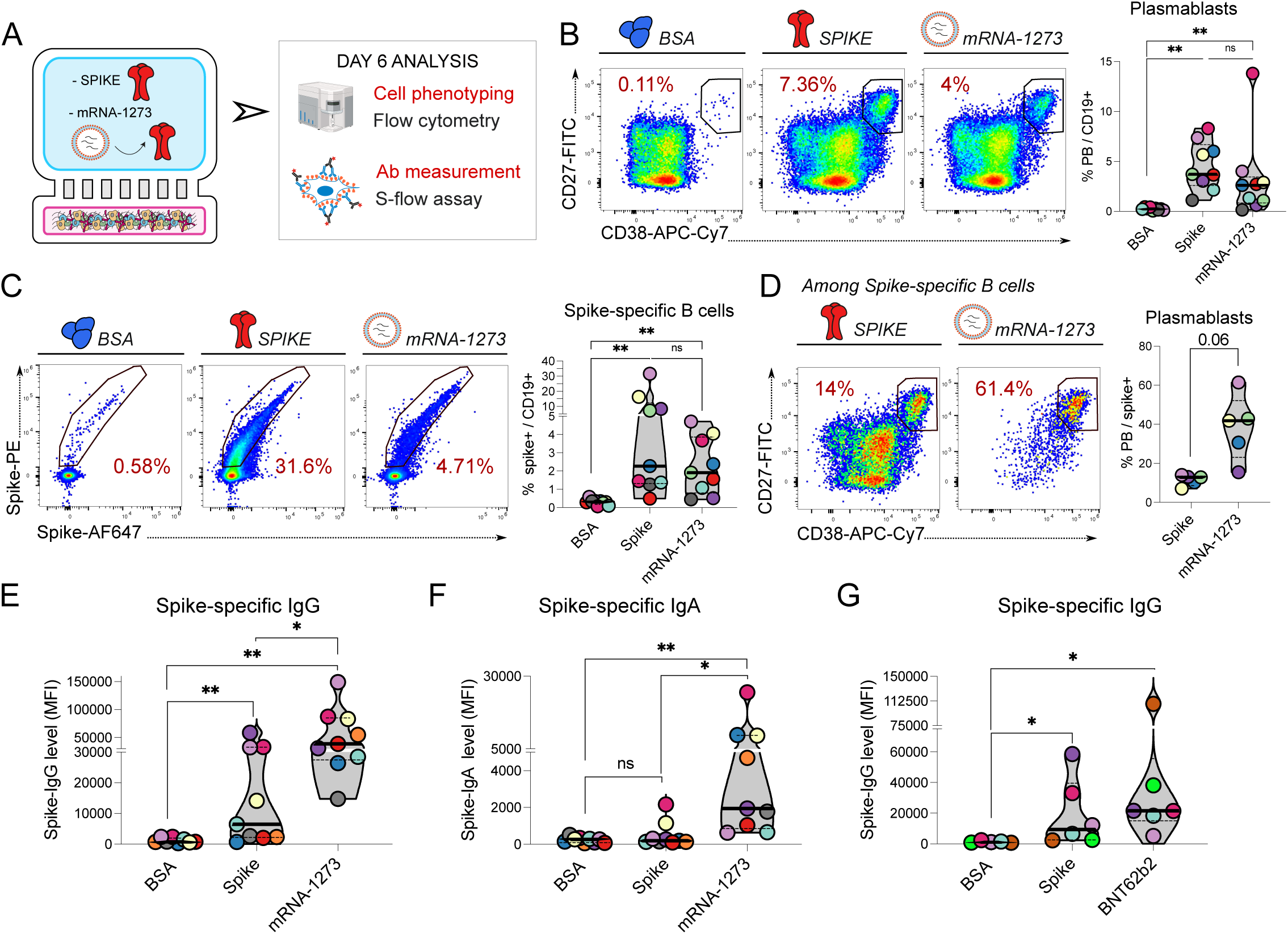
Perfusion of an mRNA vaccine induces Spike-specific B cell responses in the LO chip. (A) Experimental setup: LO chips were perfused with BSA, Spike protein, or an mRNA vaccine encoding the Spike (mRNA-1273, Moderna) and were analyzed at day 6. (B) Representative analysis of CD27 and CD38 expression in CD19+ B cells (left) and frequency of CD27hiCD38hi plasmablasts among CD19+ B cells (right). (C) Representative analysis of fluorescent Spike labeling in B cells (left) and frequency of Spike-specific cells among B cells (right). (D) Representative analysis of CD27 and CD38 expression in Spike-specific B cells (left) and frequency of CD27hiCD38hi plasmablasts among Spike-specific B cells (right). (E-F) S-flow analysis showing the mean fluorescent intensity (MFI) of Spike-specific IgG (E) and IgA (F) bound to Spike-expressing 293T cells. (G) The LO chips were perfused for 6 days with BSA, Spike protein, or an mRNA vaccine encoding the Spike (BNT62b2, Pfizer). (B-G) Each color represents an independent donor. (B-G) Differences were evaluated with a Wilcoxon matched pairs test; *p <0.05; **p < 0.01; ns: not significant.

### The LO chip captures individual variations in response to different mRNA vaccines

The SARS-CoV-2 variant that emerged in late 2021, Omicron BA.1 (or B.1.1529), proved highly divergent from previous variants, with ≥ 32 non-synonymous mutations in the Spike. Consequently, Omicron BA.1 was poorly neutralized by sera from individuals infected by previous variants and/or vaccinated against the ancestral Wuhan strain (*44*). To address this public health challenge, bivalent mRNA vaccines encoding both Wuhan and Omicron Spikes were engineered and rapidly deployed worldwide (*45–47*). We set to evaluate the *in vitro* boosting capacity of such a bivalent vaccine, by testing B cell responses induced in the LO chip by the mRNA-1273.214 vaccine (Moderna), made of LNP containing equal amounts of Wuhan and Omicron BA.1 Spike mRNA. We generated LO chips with PBMC from 7 volunteers who donated blood in 2023 and compared their in-chip B cell responses to stimulation with equivalent doses of mRNA-1273 and mRNA-1273.214 (corresponding to 3 µg mRNA coding for Wuhan S or 1.5 µg + 1.5 µg mRNA coding for Wuhan S and BA.1 S, respectively). For these experiments, the LO chips were cultured for 14 days rather than 6 days, to increase the rate of neutralizing antibody detection (Fig. 8A).

**Figure 8.**
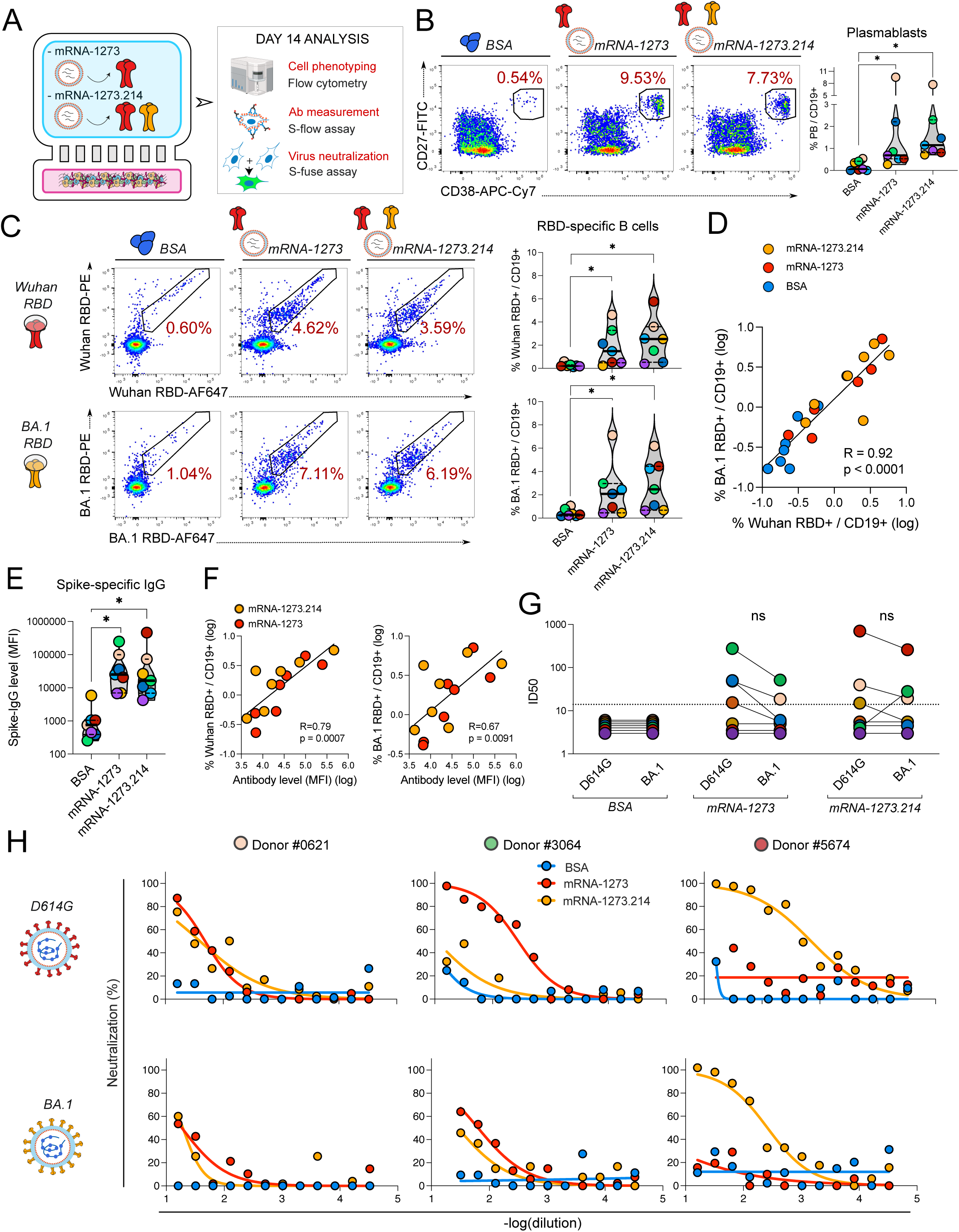
Evidence for immunological imprinting after bivalent mRNA vaccine boosting in the LO chip. (A) Experimental setup: LO chips were perfused for 6 days with BSA, a monovalent mRNA vaccine encoding the Wuhan Spike (mRNA-1273, Moderna), or a bivalent mRNA vaccine encoding the Wuhan and Omicron BA.1 Spikes (mRNA-1273.214, Moderna). (B) Representative analysis of CD27 and CD38 expression in CD19+ B cells (left) and frequency of CD27hiCD38hi plasmablasts (PB) among B cells (right). (C) Representative analysis of CD19+B cells binding to the receptor binding domain (RBD) of the Wuhan strain (top row) or the BA.1 strain (bottom row) (left) and frequency of Wuhan (top) and BA.1 (bottom) RBD-specific cells among B cells (right). (D) Positive correlation between the frequency of Wuhan RBD-specific B cells and the frequency of BA.1 RBD-specific B cells; the linear correlation coefficient R and the associated P value are reported. (E) S-flow assay analysis showing the mean fluorescent intensity (MFI) of Spike-specific IgG associated bound to Spike-expressing 293T cells. (F) Positive correlation between Spike-specific IgG levels and the frequency of Wuhan (left) or BA.1 (right) RBD-specific B cells; the linear correlation coefficient R and the associated P value are reported. (G) The neutralization capacity of antibodies produced in the LO chip was assessed by the S-Fuse assay. The threshold for the inhibitory dilution 50 (ID50) was set as 14 based on negative prepandemic controls (dashed line). ID50 are reported for neutralization of the ancestral-like D614G strain and the Omicron BA.1 strain, after stimulation with BSA, mRNA-1273, and mRNA-1273.214. (H) The dose response of D614G neutralization (top) and BA.1 neutralization (bottom) is reported in function of chip extract dilution for three representative donors to illustrate individual variability. (B, C, E, G) Each color represents an independent donor. Differences were evaluated with a Wilcoxon matched pairs test; *p <0.05; **p < 0.01; ns: not significant.

B cell maturation was induced by both mRNA vaccines to equivalent levels, as measured by the percentage of PB that persisted at day 14 (Fig. 8B). To measure specific B cells, we focused the analysis on cells that could bind the receptor binding domain (RBD) of the Spike, as this highly variable domain better distinguishes Wuhan and BA.1 specificities, while remaining critical for SARS-CoV-2 neutralization (*48*). After 14 days of culture, the monovalent mRNA-1273 and the bivalent mRNA1273.214 vaccines induced an equivalent percentage of B cells specific to the Wuhan RBD (Fig. 8C, top row; medians of 1.49% and 2.53% of Wuhan RBD+ cells, respectively). More surprisingly, the monovalent and bivalent vaccines also induced an equivalent percentage of B cells specific to the BA.1 RBD (Fig. 8C, bottom row; medians of 2.08% and 2.47% of BA.1 RBD+ cells, respectively). There were individual instances where the bivalent vaccine induced a higher amplification of RBD-specific B cells than the monovalent vaccine (for instance, donor #5674 represented by a red dot), though the reverse situation could also be observed. In all cases, the pattern observed by measuring Wuhan RBD-specific cells (top row) was similar to that observed by measuring BA.1 RBD-specific cells (bottom row). Further analysis confirmed that measurements of B cells specific for the Wuhan and BA.1 RBD showed a strong positive correlation (Fig. 8D, R=0.92, P<0.0001). These findings suggested that most of the specific B cells induced by monovalent or bivalent mRNA vaccine boosting in the LO chip cross-reacted with both RBD, consistent with the notion of immune imprinting (*48*).

We next analyzed the levels of antibodies induced in the LO chip by the S-flow assay and observed the induction of equivalent amounts of Spike-specific IgG after a monovalent or bivalent vaccine boost (Fig. 8E). The levels of antibodies detected correlated positively with the frequencies of B cells specific to the Wuhan RBD (R=0.79, P<0.001) and to the BA.1 RBD (R=0.67, P<0.01) (Fig. 8F), consistent with a direct production of antibodies by the specific B cells amplified in the chips upon boosting. We then evaluated the neutralizing capacity of antibodies produced in the chips towards the SARS-CoV-2 D614G variant, which is very close to the ancestral Wuhan strain (*49*), and the Omicron BA.1 variant. To this goal, we used the S-Fuse assay, which measures the capacity of antibodies to inhibit the fusion of reporter cells infected with the variants of interest. This assay has been previously validated for measuring neutralization capacity towards a wide array of SARS-CoV-2 variants (*44*, *50*). After mRNA-1273 boosting, antibodies neutralizing the D614G strain could be detected in 4 out of 7 donors (Fig. 8G, middle), as defined by a half-maximal inhibitory dilution above the threshold value (ID50>14). These 4 donors showed a trend for lower neutralization of BA.1, consistent with a boosting of Wuhan-specific responses. After bivalent mRNA-1273.214 boosting, antibodies neutralizing D614G could be detected in 3 out of 7 donors (Fig. 8G, right). There was no consistent increase in BA.1 neutralization, even though the bivalent vaccine encoded the BA.1 Spike. Overall, boosting with the bivalent vaccine did not confer a significant advantage over the monovalent vaccine in inducing BA.1-specific B cells or BA.1 neutralizing antibodies. These findings are in line with results from *in vivo* vaccination trials pointing to the limited efficacy of a bivalent boost at inducing Omicron-specific B cell responses (*48*, *51*, *52*).

Examination of individual neutralization curves showed various case scenarios (Fig. 8H). Some donors showed equivalent responses to monovalent and bivalent boosts (left; compare red and orange curves); other donors showed a better response to the monovalent vaccine (middle), pointing to a bias towards ancestral strain antigens; and one donor showed a better response to the bivalent vaccine (right), suggesting a recent exposure to Omicron antigens. The LO chip system could thus capture individual variability in responses to an mRNA vaccine boost, reflecting the complexity of the immunological landscape in a given population.

## DISCUSSION

We developed a microfluidic-based system that recapitulates several important features of a recall B cell response in lymphoid tissue, including a massive amplification of Ag-specific memory B cells, a concomitant induction of a specific CD4+ T cell response, the spontaneous formation of cellular clusters enriched in CD4+ T cell/B cell interactions, the induction of Ig class switch recombination, the maturation of PB with a high affinity for the stimulating Ag, and the spontaneous emigration of these PB from the lymphoid tissue-like compartment. Further, Spike stimulation in the LO chip induced the production of specific antibodies in sufficient amounts to evaluate their neutralizing capacity, a key point in developing a preclinical system for vaccine evaluation. The LO chip was designed to include the myeloid cell population naturally present in PBMC, which enabled the capture and expression of mRNA-LNP vaccines. Proof of concept experiments showed that a single bivalent mRNA vaccine boost in the chip was insufficient to shift the specificity of neutralizing antibodies towards the Omicron variant, consistent with *in vivo* vaccination trials. To our knowledge, the LO chip model represents the first preclinical model suitable for the evaluation of human B cell responses upon mRNA vaccine boosting. As setting-up an LO chip requires only a 10 million human PBMC sample, this system should prove useful to evaluate responses to candidate booster vaccines in a variety of populations that differ in immune competency and infection or vaccination history.

Comparing perfused LO chips to 3D static cultures highlighted the importance of a continuous fluid flow for the maturation of B cell responses. PB differentiation and antibody production occurred early in the LO chip after Ag boosting (day 6), with a kinetics similar to that observed in vivo (*2*). In contrast, PB showed only a limited increase, and specific antibodies were undetectable in static 2D and 3D cultures. CD4+ T cell responses appeared less dependent on fluid flow, as cytokine producing Spike-specific CD4+ T cells were significantly increased in the static 3D gel culture, though at lower levels than in the LO chip. The particularly high metabolic demands on PB that ramp up antibody secretion (*53*) may make these cells more dependent on the continuous supply of nutrients and oxygen provided by microfluidic devices. Another factor promoting B cell maturation may be the continuous perfusion of antigen, as studies in animal models show that slow antigen delivery over several days is superior to bolus antigen injection to sustain GC activity and antibody maturation (*54*). The optimized environment provided in the microfluidic chip may help explain why we could detect antibody production as early as day 6, while 9-12 days are usually required in static 2D cultures, which in addition require stimulation with superantigens rather than classic Ag (*8*, *9*). It was noteworthy that PB differentiated in the LO chip bound the Spike Ag more efficiently than activated memory B cells from the same chip, as measured by the intensity of flow cytometry labeling. This higher Ag binding capacity could not be accounted for by an avidity effect, as IgG/IgM surface expression was decreased rather than increased in PB. This suggested that the higher Ag binding capacity of PB resulted from a higher affinity of their surface Ig, which mimicked the enhanced affinity of PB that differentiate within GC *in vivo* (*1*, *3*). The fact that PB differentiated in the LO chip preferentially egressed from the tissue-like compartment to join the perfused compartment also recapitulated an important aspect of the maturation of the B cell response *in vivo* (*10*). The migratory behavior of PB in the LO chip may also have practical applications for monoclonal antibody production, as cells harvested from the upper chip compartment are highly enriched in high-affinity Ag-specific B cells, which should facilitate the cloning of BCR of interest.

A CD4+ T cell/B cell dialogue was established in the LO chip upon Ag perfusion, as indicated by the concomitant induction of CD4+ T cell and B cell specific responses, and the positive correlation between PB induction and ICOS+ CD38+ CD4+ T cell frequencies. The spontaneous organization of CD4+ T cell/B cell clusters upon Ag perfusion highlighted a direct interaction between the two cell types, with tight contacts suggestive of immunological synapse formation. Consistent with this notion, the CD4+ T cells found within the clusters had an activated phenotype, based on large size and ameboid morphology, and expressed the costimulatory molecule ICOS, a marker upregulated in Tfh cells (*55*) and involved in driving B cell maturation (*56*). The CD4+ T cells within the clusters were thus likely engaged in providing help to B cells, similar to their function in GC. The induction of CXCL13 chemokine expression, a well-established marker of GC activity (*37*), further supports this notion. It should be noted that we did not detect a segregation between a light zone enriched in activated CD4+ T cells and a dark zone enriched in proliferating B cells within the cell clusters. Therefore, the LO chip recapitulates some important features of GC, but not all of them, and may be further improved, for instance by the inclusion of stromal cells that would structure lymphoid tissue territories through the secretion of chemokines (*10*). The current LO chip model retains the advantage of providing CD4+ T cell help in an Ag-specific fashion, in contrast to systems requiring superantigen stimulation, CD40L-expressing fibroblasts, or exogenous cytokine addition (*15*, *16*, *27*). This may be one reason why immune response analysis in LO chips was not hampered by spontaneous CD4+ T cell or B cell activation, as shown by low/undetectable responses upon stimulation with the irrelevant BSA Ag. High-density cultures of purified human T and B lymphocytes are known to be prone to spontaneous activation (*28*), which can be an asset to predict the capacity of biologics to generate a cytokine storm (*57*) but is a hindrance in the study of Ag-specific responses. The LO chip design includes diverse leukocyte subsets, such as NK cells that can immunoregulate T and B cell populations (*58*), possibly accounting for the lack of spontaneous T/B cell activation. The LO chip thus appears well-suited to the combined evaluation of Ag-specific T and B memory responses from human donors.

We could also monitor the activation of myeloid cells in LO chips. By harvesting the chips at day 2, we could document the early induction of activation (HLA-DR) and costimulation (CD40, CD86) markers on DC and monocytes after mRNA vaccine treatment. Labeling with an anti-Spike antibody showed these two cell types were primarily responsible for expressing the mRNA-encoded Spike, consistent with *in vitro* and *in vivo* studies of mRNA-LNP delivery (*41*, *59*). The early activation of these APC was compatible with the notion that mRNA-LNP particles possess an intrinsic adjuvant effect, due to the ionizable or PEG-conjugated lipids forming the particle envelope and to their nucleic acid content (*38*, *43*). Interestingly, monocytes in mRNA-1273 treated chips showed signs of differentiation towards a DC phenotype, as indicated by the induction of the CD11c and DC-SIGN markers. This suggested that mRNA-LNP treatment could promote a myeloid differentiation pathway conducive to antigen presentation. The preferential upregulation of HLA-DR and costimulatory markers in myeloid cells that expressed the Spike after mRNA-1273 treatment also supported the idea of an improved antigen presentation capacity in cells that had captured the vaccine. The presence of PBMC-derived myeloid cells and their differentiation into functional APC represents an asset of the LO chip system, as it enables the evaluation of new generation mRNA vaccines, in contrast to systems that rely on purified B cells and CD4+ T cells. The presence of other leukocyte subsets may also be relevant, and their role remains to be explored. For instance, a recent report suggests that NK cells may limit the expression of mRNA vaccines by killing Spike expressing cells through an Fc-dependent mechanism, if Spike-specific antibodies preexist in the circulation (*60*). The LO chip system opens the possibility of modulating leukocyte populations and circulating antibody concentrations to better model parameters involved in the efficiency of recall responses to mRNA vaccines.

The fact that we detected strong Spike-specific antibody production in mRNA-1273 treated samples suggests that the entire chain of events needed for B cell maturation takes place in the LO chip, including mRNA-LNP capture by myeloid cells, release of the mRNA from endosomes into the cytosol, mRNA translation, expression of the resulting Spike protein at the myeloid cell surface, and capture of this protein by cognate B cells, leading to BCR activation. Further, the formation of B cell/CD4+ T cell clusters suggests that B cells can internalize, process, and present Spike epitopes at their surface, enabling the provision of help by CD4+ T cells specific for these epitopes, and promoting B cell maturation into antibody secreting PB. The production not only of Spike-specific IgG but also of Spike-specific IgA suggests that mRNA vaccine boosting promotes immunoglobulin class-switching in the LO chip. Further, the higher frequency of PB among Spike-specific B cells induced after an mRNA vaccine boost compared to a Spike protein boost supports the idea that mRNA vaccines are self-adjuvanting and trigger rapid memory B cell maturation (*61*). The fact that the mRNA vaccine induces expression of Spike proteins anchored at the plasma membrane may also promote the crosslinking of multiple BCRs at the surface of specific B cells, leading to a more potent maturation signal. Thus, both the self-adjuvanting properties of mRNA-LNP and their capacity to encode proteins in their native conformation may contribute to the success of this emergent vaccination platform (*38*).

Immune escape is recognized as a key parameter driving the successive waves of SARS-CoV-2 variants worldwide (*62*, *63*), spurring the development of variant-specific booster vaccines. The worldwide spread of the highly divergent Omicron variant starting from late 2021 led to the development of bivalent mRNA vaccines encoding the ancestral Wuhan Spike and an Omicron BA.1 or BA4/5 Spike, matching the variants circulating in early and late 2022, respectively (*64*). The bivalent vaccines were rapidly deployed and could boost the induction of SARS-CoV-2 neutralizing antibodies, ensuring a protection from severe COVID (*46*, *65*). However, it soon emerged that, in most studies, the titers of antibodies specific to Omicron were not preferentially increased by the bivalent vaccine as compared to those induced by the monovalent Wuhan vaccine (*66*, *67*). Previous studies had also reported a lower induction of Omicron-neutralizing antibodies after breakthrough Omicron infection in individuals who had been vaccinated multiple times against the Wuhan strain compared to unvaccinated individuals (*31*, *68*). There is now convergent evidence that immune imprinting is limiting the induction and efficiency of Omicron-specific responses in individuals with preexisting B cell memory to previous SARS-CoV-2 variants, a phenomenon that can be explained by the preferential amplification of preexisting memory B cells upon vaccine boosting (*48*, *51*, *69*)(*70*).

Given these complex immunological interactions, we set out to model monovalent and bivalent mRNA vaccine boosting in the LO chip. The comparison of response induced by the original Wuhan vaccine mRNA-1273 and the bivalent Wuhan/BA.1 vaccine mRNA-1273-214 revealed only moderate differences between the two formulations, with an equivalent frequency of PB induction and an equivalently high level of Spike-specific IgG production. A limitation of the study was that only IgG specific for the Wuhan strain were detected in the S-flow assay. However, we were able to use strain-specific reagents to measure the frequency of RBD-specific B cells and found a comparable amplification of B cells specific for the Wuhan RBD by the two vaccines, but also, less expectedly, a comparable induction of B cells specific for the BA.1 RBD. The tight correlation between the frequencies of B cells specific for the Wuhan and BA.1 RBD (R=0.92, P<0.0001) suggested that most of these B cells were cross-reactive to the two strains. Thus, the LO chip system appeared to recapitulate the phenomenon of immune imprinting observed *in vivo*, with a preferential expansion of cross-reactive B cells upon vaccine boosting. Recent findings suggest that repeated boosting with an Omicron-derived monovalent mRNA vaccine can overcome immune imprinting by the ancestral strain (*52*), a notion that could be tested in the LO chip in future studies.

The production of Spike-specific antibodies at day 14 in the LO chip proved sufficient to measure their neutralizing capacity, which is an asset of the LO chip system, by opening the possibility of an i*n vitro* evaluation of vaccine efficacy. The induction of SARS-CoV-2 neutralizing antibodies could be detected in 6 out of 7 donors tested, though with different individual patterns. The monovalent vaccine tended to induce higher neutralizing antibody levels against the ancestral D614G strain than the BA.1 strain. In contrast, the bivalent vaccine did not induce a higher neutralization of BA.1 than of D614G. Further, comparing the two vaccines, we did not detect an increased neutralization of BA.1 after bivalent boosting compared to monovalent boosting, compatible with *in vivo* data (*66*, *67*), and reinforcing the notion that immune imprinting limited the emergence of Omicron-specific neutralizing antibodies. Examination of individual neutralization curves revealed different case scenarios, with donors responding equivalently well to the two vaccines, donors responding more efficiently to the monovalent vaccine, suggestive of strong imprinting, and one donor responding better to the bivalent vaccine, possibly due to a recent infection with an Omicron-derived variant. This analysis illustrates the diversity of immunological histories in the population, and the resulting individual variability in vaccine responses. In the face of such variability, the LO chip can provide a useful preclinical system to evaluate the capacity of candidate vaccines to induce neutralizing antibodies against current SARS-CoV-2 variants in diverse human populations.

In conclusion, we developed a versatile Lymphoid Organ-Chip model suitable for the preclinical evaluation of human recall responses to different vaccine formulations, including the new generation mRNA vaccines. The LO chip recapitulates the early activation of myeloid cells upon antigen capture, and the later activation of antigen specific CD4+ T cell and B cells, thus capturing the multifaceted aspects of a recall response. The formation of CD4+ T cell/B cell clusters and the differentiation and emigration of plasmablasts mimicked important features of B cell maturation within lymphoid tissues. Further, the microfluidic perfusion promoted a massive amplification of antigen-specific memory B cells and plasmablasts, enabling the detection of secreted antibodies and the evaluation of their neutralizing capacity. This approach confirmed the efficiency of mRNA vaccines at boosting antibody responses, but also the lack of advantage of a bivalent over a monovalent vaccine boost. The LO chip thus represents a streamlined 3D organ model applicable to the evaluation of vaccine boosters in diverse human populations, which should be an asset in the face of a rapidly evolving SARS-CoV-2 pandemic.

## MATERIALS AND METHODS

### Human blood samples

Healthy donors were anonymous volunteers who donated blood at the Etablissement Français du Sang (EFS). Blood samples were provided to Institut Pasteur (Paris) under agreement C CPSL UNT - N°18/EFS/041. PBMC were isolated from buffy coats or cytapheresis collars via density gradient centrifugation on lymphocyte separation medium (Eurobio, #CMSMSL01-01) and were resuspended in fetal bovine serum (FBS; PAN Biotech, #P30-3306) containing 10% dimethyl sulfoxide (DMSO) (Sigma Aldrich, #226827) before cryopreservation in liquid nitrogen. For culture, PBMC were thawed and resuspended in complete medium consisting in RPMI-1640 glutaMAX ^TM^ (Gibco, #61870036) supplemented with 100 U/mL penicillin/streptomycin (Gibco, #15140122), 10 mM HEPES (Gibco #15630056), 1 mM sodium pyruvate (Gibco, #15630056), 0.1 mM non-essential amino acids (Gibco, #11140050,), 0.05 mM 2-mercaptoethanol (Gibco, #31350010), and 10% heat-inactivated human AB serum (Institut de Biotechnologies, #201021334). PBMC were rested for a minimum of two hours in complete medium supplemented with 1 µg/mL of DNAse from bovine pancreas (ThermoFischerScientific, #58409400) prior to culture. Plasma collected during PBMC preparation was used to screen donors for the presence of SARS-CoV-2 Spike-specific IgG, using the S-flow assay (see below). Only the donors seropositive for SARS-CoV-2 were included into the study.

### Microfluidic chip activation and preparation for cell seeding

Two-compartment S1^®^ chips from Emulate (Boston, MA) were used throughout the study. S1 chips are composed of polydimethylsiloxane (PDMS), a transparent and gas permeable elastomeric polymer, and consist in two parallel channels separated by a membrane with 7 µm pores. Fluid flow can be controlled independently in each channel through a Zoe^®^ culture module connected to an Orb^®^ hub module (Emulate), with continuous medium perfusion ensuring the renewal of essential nutrients and enabling primary cell culture at high density. Components secreted by cultured cells can be collected in the two reservoirs connected to the outlet of each channel and contained within each chip holder, or Pod^®^ (Emulate). Prior to culture, the inner PDMS surface of the chip channels were activated according to the manufacturer guidelines to enable proper ECM attachment. The chips were then coated at 4°C overnight with a solution of 30 µg/mL of type I collagen (Corning, #354236) and 100 µg/mL Matrigel (Corning, #354234) diluted in phosphate buffered saline (PBS) (Gibco, #14190-094). Before cell seeding, the chips were incubated for at least 1 hour in a culture incubator at 37°C, and each channel was then washed twice with 200 µL of complete medium. Just prior to cell seeding, the medium in each channel was thoroughly aspirated.

### Lymphoid Organ-chip (LO chip) culture

The LO chip design consists in a high-density 3D human PBMC culture within an ECM gel in the lower channel of an S1 chip, while medium and antigens are fluxed in the upper channel of the chip. For LO chip seeding, PBMC were resuspended at the concentration of 580 million cells/mL in an ECM composed of half collagen I solution and half degassed complete medium, to achieve a final concentration of 1.5 mg/mL of type I collagen. The initial type I collagen solution at 3 mg/mL was prepared by mixing rat tail type I collagen (Corning, #354236), sterile water, DPBS 10X, and adequate amount of 1N hydroxide sodium to reach pH = 7.4 (Sigma Aldrich, #655104), allowing gelation. The complete medium was degassed using aspiration in a Steriflip-HV cup (Merck, #SE1M003M00). All reagents and PBMC were maintained on ice before seeding. 17 µL of the viscous cell/ECM solution containing a total of 10 million PBMC was then introduced through the inlet port to fill the lower channel of the chip. The cell/ECM solution was allowed to gel for 30 min in a cell culture incubator at 37°C and 5% CO2. After gelation, a thin disc-shaped plastic membrane was inserted to block the inlet port of the gel-filled channel, as a measure to prevent bubble formation within the ECM. The disc forms a fluid- and air-tight barrier between the chip inlet port and Pod^TM^ inlet reservoir, while still allowing pressure to be applied via the chip lower channel outlet. The outlet reservoir was filled with 1 mL of complete medium to avoid drying of the ECM. The chips were perfused through the upper channel inlet with 30 µL/h of complete medium, as programmed on the Orb^TM^ controller (Emulate), with the chip culture module placed in an incubator maintained at 37°C and 5% CO2. At day 0, the inlet reservoir connected to the upper channel was filled with 3 mL of complete medium supplemented with antigen. Every 3 days, the perfused medium was renewed with a 1:1 mix of effluent medium and fresh complete medium. The inclusion of effluent medium ensured maintenance of the cytokine milieu induced by antigenic stimulation and continuous perfusion of the antigen.

### Antigens used to stimulate LO chips

The protein antigens tested included the control protein bovine serum albumin (BSA, Invitrogen, #AM2616) at 1 µg/mL and the Spike protein of the SARS-CoV-2 Wuhan strain (BEI Resources, #NR-53937) at 1 µg/mL. The mRNA-based lipid nanoparticle (LNP) vaccines used included the monovalent Wuhan vaccine mRNA-1273 (Moderna), the bivalent Wuhan/Omicron BA.1 vaccine mRNA-1273.214 (Moderna); and the monovalent Wuhan vaccine BTN162b2 (Pfizer), all tested at 3 µg/mL. Control LNP were engineered with a “Moderna-like” lipid composition by the Oz Biosciences company and contained mRNA encoding for the control protein ovalbumin (OVA-LNP) or for the green fluorescent protein (GFP-LNP). Control LNP were used at a 3µg/mL concentration, similar to the one used for mRNA vaccines.

### Static 2D and 3D culture**s**

2D or 3D classical cultures were performed to evaluate antigenic responses in conditions devoid of fluid flow. For static 2D cultures, 10 million PBMC were resuspended in 1 mL of complete medium in a 24-well plate. For static 3D cultures, 10 million PBMC were resuspended in an ECM droplet at the same concentration used in the LO chip (10 million cells/17µL), and the cell/ECM gel droplet was deposited in 1 mL of complete medium in a 24-well plate. For stimulation, antigen (BSA, Spike, or mRNA vaccine) was added at the same concentration as the one used in the LO chip.

### Phenotyping of immune cell subsets

At each timepoint (day 2, or day 6, or day 14), effluent from the upper outlet reservoir was collected, aliquoted, and stored at -20°C. Chips were disconnected from the Pods, and upper channels were washed twice with 100 µL PBS introduced in the inlet port, while the outlet port was connected to an empty 200 µl tip. The collected cells were either pooled with those recovered from the lower channel or analyzed separately to characterize cell migration from the lower to the upper channel. To harvest cells from the lower channel, the ECM had first to be digested. To do so, all the chip ports were connected with empty tips, except the upper inlet port which was manually perfused with 100 µL of Cell Recovery Medium (Corning, #354253) or of Cultrex Organoid Digestion solution (R&D systems, #3700-100-01). Chips were incubated for 45 minutes at 4°C to allow diffusion of the digestion solution from the upper to the lower channel. After the incubation, the lower inlet port was connected to a new tip with 80 µL of cold PBS that was manually perfused through the lower channel to flush the cells and ECM remnants out of the chip for collection. The lower channel was further washed twice with 80 µL of cold PBS, and all the washes were pooled prior to centrifugation at 300 g for 5 min. The pelleted cells were resuspended in PBS for immunostaining while the supernatants from ECM digestion were collected and stored at -20°C.

For the phenotyping of myeloid cells, the cells collected from LO chips at day 2 were resuspended in cold PBS, incubated for 10 min at 4°C with a Live/Dead fixable dye (Invitrogen, #L10119), and then stained for 30 min at 4°C in PBA buffer (PBS, 0.5% BSA, 2 mM EDTA) in the presence of Fc Block (1:50 dilution, BD-Biosciences), using the following antibody combination: CD19, CD3, CD4, CD14, DC-SIGN, CD11c, HLA-DR, CD40, CD86 and anti-Spike Ab102-AF647 (provided by Cyril Planchais and Hugo Mouquet, Institut Pasteur) (*42*). Cells were then washed twice in PBS and fixed with a Cytofix/cytoperm kit (BD Biosciences, 554655) for 20 min at 4°C. Intracellular staining for anti-Spike Ab102 was done for 30 minutes at 4°C, to detect intracellular Spike. The irrelevant mGO53 human IgG1 was used as an isotypic control. After staining, cells were washed twice in PBS, fixed in a 4% paraformaldehyde (PFA) in PBS solution for 20 min at room temperature (RT), and resuspended in cold PBS.

For detection of Spike specific or RBD-specific B cells, recombinant biotinylated Spike (Miltenyi: Wuhan spike #130-127-682; Omicron BA.1 spike1#30-130-417) or biotinylated RBD (Miltenyi: Wuhan RBD #130-129-570; Omicron BA.1 RBD #130-130-419) were first coupled with AF647-streptavidin (Invitrogen, S32357) and PE-streptavidin (Miltenyi, 130-106-790) at a 5:1 molar ratio in cold PBA buffer for a minimum of 15 min. PBMC were stained with the fluorescent RBD or Spike at 3 µg/mLfor 45 min in cold PBA buffer. After two PBS washes, cells were incubated for 10 min at 4°C with a Live/Dead fixable dye (Invitrogen, #L34963) in PBS, and then stained in PBA with Fc Block (1:50 dilution) using the following combination of antibodies: CD3, CD4, CD19, CD27, CD38, IgG and IgM. Cells were washed twice in PBS, fixed in a 4% paraformaldehyde in PBS solution, and resuspended in cold PBS.

For detection of spike-specific CD4+ T cells, the harvested cells were restimulated using a pool of overlapping Wuhan Spike peptides (Miltenyi, 130-126-701) at 2 µg/mL for 16 hours in the presence of brefeldin A (Biolegend, #400602) at 5 µg/mL. To detect cytokine secretion by CD4+ T cells, cells were resuspended in cold PBS, incubated with a Live/Dead fixable dye (Invitrogen, #L10119) for 10 min at 4°C, and surface stained for CD3 and CD4 in PBA buffer with Fc block (1:50 dilution) for 20 min at 4°C. Cells were washed twice and then fixed and permeabilized with a Cytofix/Cytoperm kit (BD Biosciences, 554655) for 20 min at 4°C. Intracellular staining for IFN-γ, TNF-α, and IL-2 was done for 30 minutes at 4°C. Cells were centrifuged and resuspended in Fix/Perm buffer and stored at 4°C prior to cytometry analysis.

Flow cytometry analysis was carried out using an Attune NxT flow cytometer (ThermoFisherScientific) and data were analyzed using the FlowJo V10 software (Flowjo, LLC). The antibody references and dilution used are listed in supplementary Table S7.

### Measurement of CXCL13 production

The chemokine CXCL13 was measured in LO chip effluent using a human CXCL13 ELISA assay (Quantikine, R&D Systems #DY801), following the manufacturer’s instructions.

### S-flow antibody assay

IgG and IgA antibodies specific to the SARS-CoV-2 Spike were detected by the S-flow assay, which measures antibody binding to Spike-expressing HEK 293T cells, as previously described. This assay was shown to have a 100% specificity (95% confidence interval [CI]: 98.5%–100%) and 99.2% sensitivity (95% CI: 97.69%–99.78%) for COVID-19 patient sera, and to outperform ELISA assays in terms of sensitivity (60).The spike-expressing cells (293T-S) were generated by transducing HEK 293T cells (ATCC® CRL-3216™) with a lentivector expressing a codon-optimized Wuhan SARS-CoV-2 Spike protein (GenBank: QHD43416.1). Control 293T cells were transduced with an empty lentivector to assess background staining. The transduced cells were selected with 2.5 µg/mL of puromycin. To perform the S-flow assay, 5 x 10^4^ 293T-S cells were plated in a 96-well round bottom plate. 50 µL of patient serum diluted 1:300 in MACS buffer (Miltenyi Biotech) or 50 µL of ECM chip extract (undiluted at day 6, or diluted 1:6 at day 14) were added to the cells, and the mix was incubated for 1 hour at 4°C. The cells were then washed in PBS and stained with a mix of secondary antibodies anti-human IgG-Fc-AF647 (1:600) and anti-human IgA-Alpha-Chain-AF488 (1:200) (Thermo Fisher Scientific) for 30 min at 4°C. Cells were fixed for 10 min in 4% PFA and were acquired on an Attune NxT flow cytometer (Life Technologies). Results were analyzed with the FlowJo V10 software. For each sample, the background signal was measured in control 293T cells lacking S and subtracted to define the specific signal. The mean fluorescence intensity (MFI) of IgG and IgA binding was reported, as it was found to provide a quantitative measurement of the levels of SARS-CoV-2 Spike specific antibodies (*32*, *33*).

### Immunofluorescence microscopy

For in-chip immunofluorescence, cells were stained by relying on antibody diffusion from the upper channel to the lower channel. The LO chips were emptied of medium and fixed by filling the upper channel through the inlet port with 4 % PFA in PBS 1x, while the other ports were connected to empty tips. The chips were incubated for 30 min at RT, before aspirating the fixative solution and washing the chips 3 times with PBS for 5 min. The upper channels were then filled with 100 µL of a solution containing primary antibodies diluted in PBS with 1% FBS, 0.1% Triton-X100, and 1:20 Fc Block (BD biosciences). The following primary antibodies were used: CD4-AF488 (BD biosciences, 557695, RPA-T4, 1/25), CD19-AF561 (eBioscience^TM^, 505-0199-42, HIB19, 1/25), ICOS (Cell Signaling Technology, rabbit 89601, D1KT2^TM^, 1/100) Ki67 (ThermoFisherScientific, rabbit MA5 14520, 1/200). After three washes with 200 µL of PBS added through the upper inlet port, the chips were incubated with the secondary antibody goat anti-Rabbit IgG-AF647 (Invitrogen, A32733, polyclonal, dilution 1:200) and the nuclear dye Hoechst (Life Technologies, H3570, 1:1000) for 4 hours at 4°C in the dark. After three PBS washes, the chips were analyzed using a spinning disc confocal microscope (Nikon, Ti2E, Yokogawa, CSU W1) using a 40x objective with a long working distance, to visualize the whole depth of the chip’s lower channel.

### S-Fuse neutralization assay

Antibody neutralization was measured by the inhibition of cell fusion in a GFP-split cell system. U2OS-ACE2 GFP1–10 and U2OS-ACE2-GFP11 cells, also termed S-Fuse cells, become GFP+ when they fuse together upon productive infection by SARS-CoV-2 (*50*). The S-Fuse cells tested negative for mycoplasma. S-Fuse cells were mixed (at a 1:1 GFP1-10 / GFP11 ratio) and plated at 8 × 10^3^ cells per well in a μClear 96-well plate (Greiner Bio-One). The tested SARS-CoV-2 strains were incubated with serially diluted chip effluent or supernatants from ECM digestion for 20 min at RT, and the mixture was then added to S-Fuse cells. 18 h later, cells were fixed with 2% PFA (Electron microscopy # 15714-S), washed in PBS, and stained with Hoechst at a 1:1000 dilution (Invitrogen, H3570). Images were acquired automatically using an Opera Phenix high-content confocal microscope (Perkin Elmer). The GFP+ area and the number of nuclei per well were quantified using the Harmony software (PerkinElmer). The percentage of neutralization was computed using the number of GFP+ syncytia as value with the following formula: 100 × (1 − (value with antibodies − value in ‘non-infected’) / (value in ‘no antibodies’ − value in ‘non-infected’)). Neutralizing activity of each sample was expressed as the half maximal inhibitory dilution (ID50). Of note, we previously validated the S-fuse assay by showing a strong correlation between neutralization titers obtained with the S-Fuse reporter assay and a pseudovirus neutralization assay (*33*, *71*).

### Statistical analyses

Statistics were computed with the GraphPad Prism v10.1.1 software. The non-parametric Mann-Whitney test and the Wilcoxon matched-pairs rank test were used to compare groups. Correlations were analyzed by simple linear regressions with the associated Pearson correlation coefficient R reported. ID50 values were obtained after non-linear curve fit using a four-parameter logistic regression model in Prism. P values lower than 0.05 were considered statistically significant. The nature of statistical tests used is reported in the figure legends.

## ACKNOWLEDGMENTS

We thank Sophie Novault, at the Flow Cytometry Platform of Institut Pasteur (IP), for advice on cytometry data acquisition; Cyril Planchais and Hugo Mouquet, at the Humoral Immunology Unit of IP, for anti-Spike antibodies; Florence Guivel-Benhassine and Françoise Porrot, at the Virus & Immunity Unit of IP, for generating viral stocks; Blanca Liliana Perlaza and Marie-Noëlle Ungeheuer, at the ICAREB biobank facility of IP, for access to human cells; Céline Fichot, at the Medical Center of IP, and Catherine Perves, at Hospital Cochin, for providing mRNA vaccines. The following reagent was obtained from BEI Resources: recombinant Spike protein from the SARS-CoV-2 Wuhan strain (#NR-53937).

## FUNDING

This work was supported by the Emulate company (contract S-RD21002; L.A.C.); the COROCHIP project funded by the Pasteur COVID-19 RP call (to L.A.C. and S. G.); The Fondation de France (PR-166156 project to L.A.C.); the Urgence COVID-19 Fundraising Campaign of Institut Pasteur (PFR7 project; L.A.C); the French agency for AIDS and emerging diseases research ANRS-MIE (ECTZ213626 project; L.A.C.); and the Institut Carnot Pasteur Microbe & Santé (to S.Go.).

## AUTHOR CONTRIBUTIONS

Conceived and designed the experiments: RJM, OS, LE, MB, SGo, LAC. Performed the experiments: RJM, DP, IS, JK, HM, CC, BFF, HD, RR, SGe. Analyzed the data: RJM, DP, LAC. Wrote the paper: RJM, LAC. All authors reviewed and approved the manuscript.

## COMPETING INTERESTS

LE is an employee of Emulate and may hold equity. MB is an employee of Hoffmann-LaRoche. The company provided support in the form of salaries and overhead for RJM, but did not have any additional role in the study design, data collection and analysis, decision to publish, or preparation of the manuscript.

## DATA AND MATERIALS AVAILABILITY

All data needed to evaluate the conclusions in the paper are present in the paper and/or the supplementary materials.

