## Supplementary Figs S1 to S6 and Table S7 for "Recapitulating memory B cell responses in a Lymphoid Organ-Chip to evaluate mRNA vaccine boosting strategies"

### SUPPLEMENTARY MATERIAL CONTENT

**Supplementary Figure S1**: B cell gating strategy.

**Supplementary Figure S2**: Antibodies concentrate into the extracellular matrix (ECM) of the LO chip.

**Supplementary Figure S3**: Spike perfusion induces spatial organization and proliferation of lymphocytes in the LO chip.

**Supplementary Figure S4**: Comparison of 2D and 3D static cultures.

**Supplementary Figure S5**: Myeloid cell gating strategy.

**Supplementary Figure S6**: Expression of a GFP-RNA lipid nanoparticle (LNP) in immune cell populations.

**Supplementary Table S7**: References of antibodies used in the study.

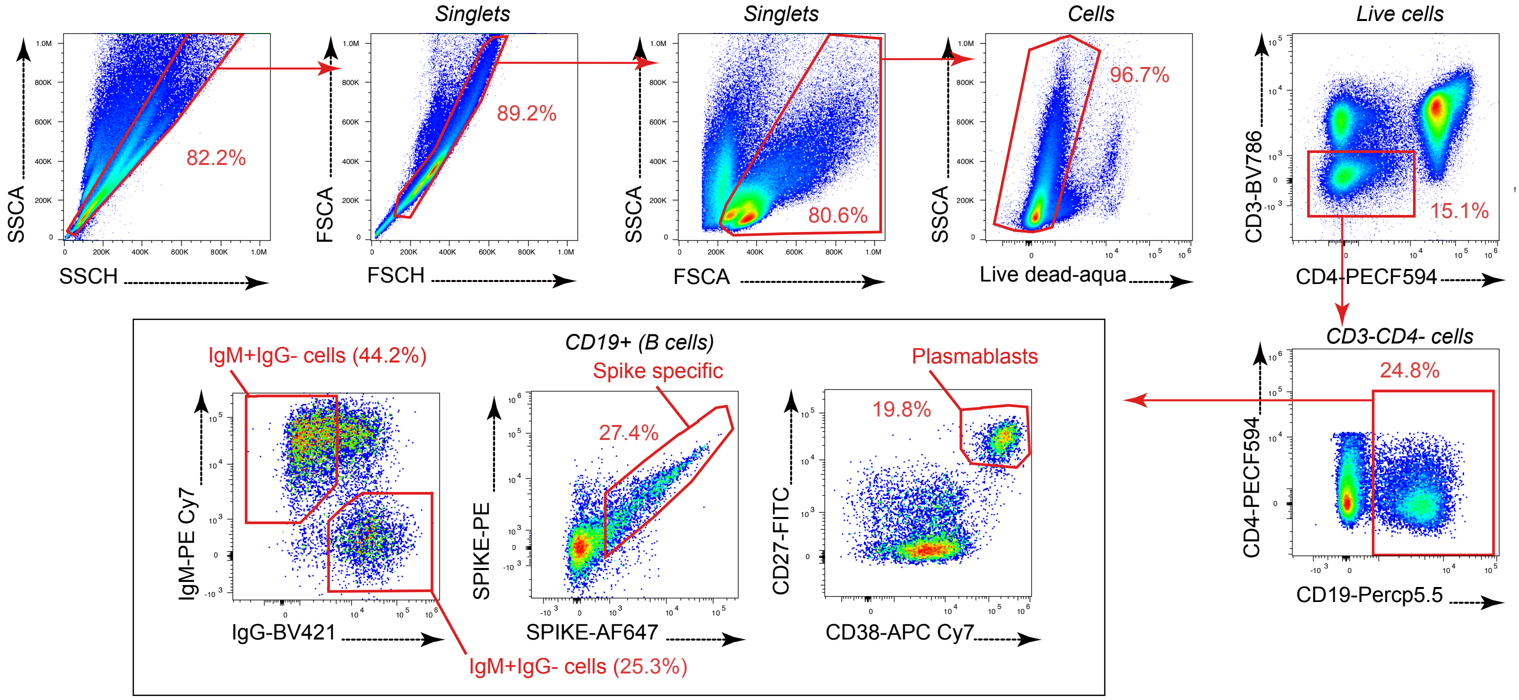


**Figure S1: B cell gating strategy.**
B cells were defined as CD3- CD4- CD19+ live singlet cells. B cell subsets were defined based on IgG and IgM expression, or on the binding of the SARS-CoV-2 Spike protein labeled with two different fluorochromes, or on the expression of CD27 and CD38, with CD27hiCD38hi cells defined as plasmablasts.

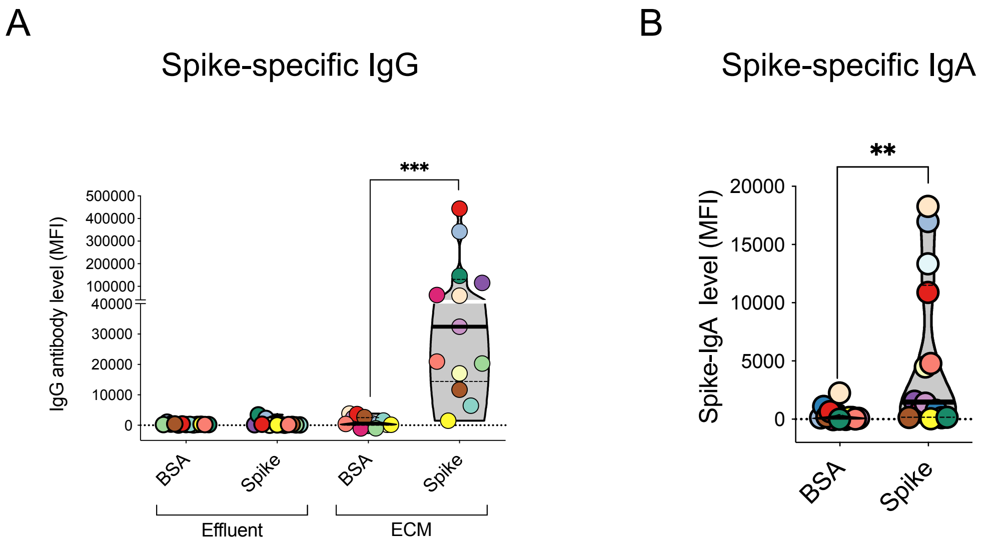


**Figure S2. Antibodies concentrate into the extracellular matrix (ECM) of the LO chip.**
(A) Antibodies were evaluated by the S-flow assay, which measures the mean fluorescent intensity (MFI) of Spike-specific IgG bound to Spike-expressing 293T cells. Spike-specific IgG were measured at day 6 in LO chip effluent medium (Effluent) and in the digestion solution obtained from the ECM recovered from the LO chip (ECM) after 6 days of stimulation with the BSA or the Spike protein. (B) Spike-specific IgA were measured by the S-flow assay in the ECM recovered from the LO chip after 6 days of stimulation with the BSA or the Spike protein. (A-B) Each color represents an independent donor. Differences were evaluated with a Wilcoxon matched pairs test *p <0.05; **p < 0.01, ***p < 0.001.

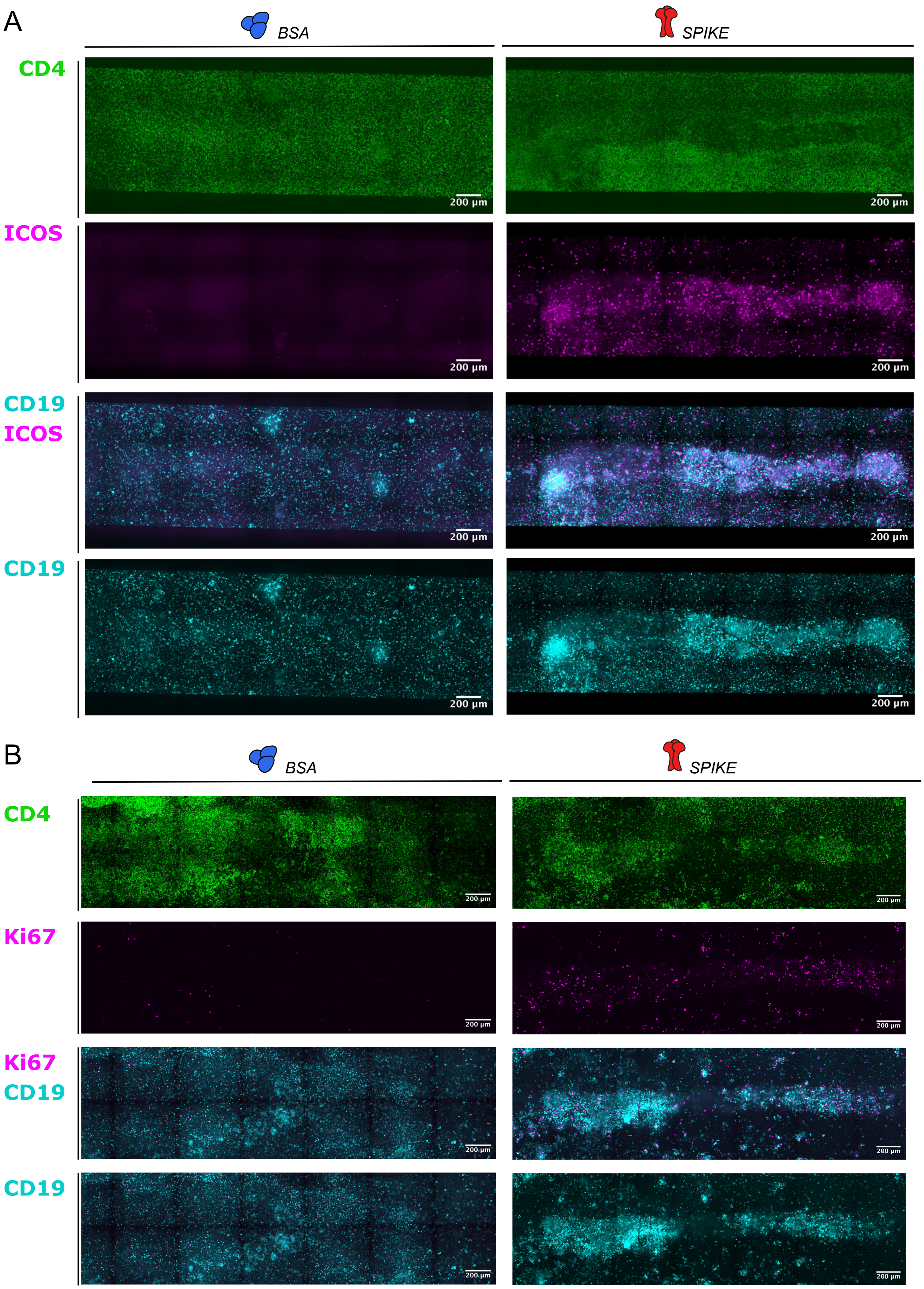


**Figure S3. Spike perfusion induces spatial organization and proliferation of lymphocytes in the LO chip.**

Confocal microscopy was used to image the lower channel of LO chips after 6 days of perfusion with the BSA or Spike proteins. Images were acquired at 20x magnification and processed with a Z stack projection on maximal intensity. (A) CD4+ T cells were labeled with CD4 (green) and ICOS (magenta) while B cells were labeled with CD19 (cyan). (B) CD4+ T cells were labeled with CD4 (green), B cells with CD19 (cyan), and proliferating cells with Ki67 (magenta).

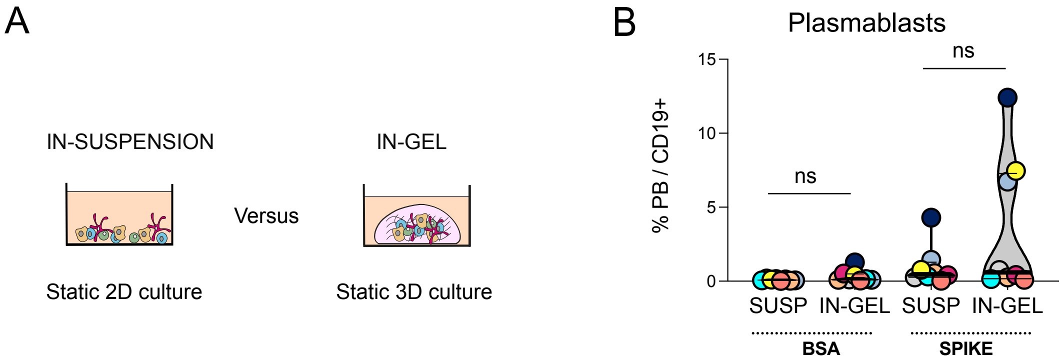


**Figure S4. Comparison of 2D and 3D static cultures.**
(A) Experimental setup: the same number of PBMC were cultivated for 6 days either in suspension or within an extracellular matrix gel droplet in a 24-well plate in the presence of the same concentration of BSA or Spike protein. (B) The frequency of CD27hiCD38hi plasmablasts (PB) among CD19+ B cells is reported. Differences were evaluated with a Wilcoxon matched pairs test. ns: not significant.

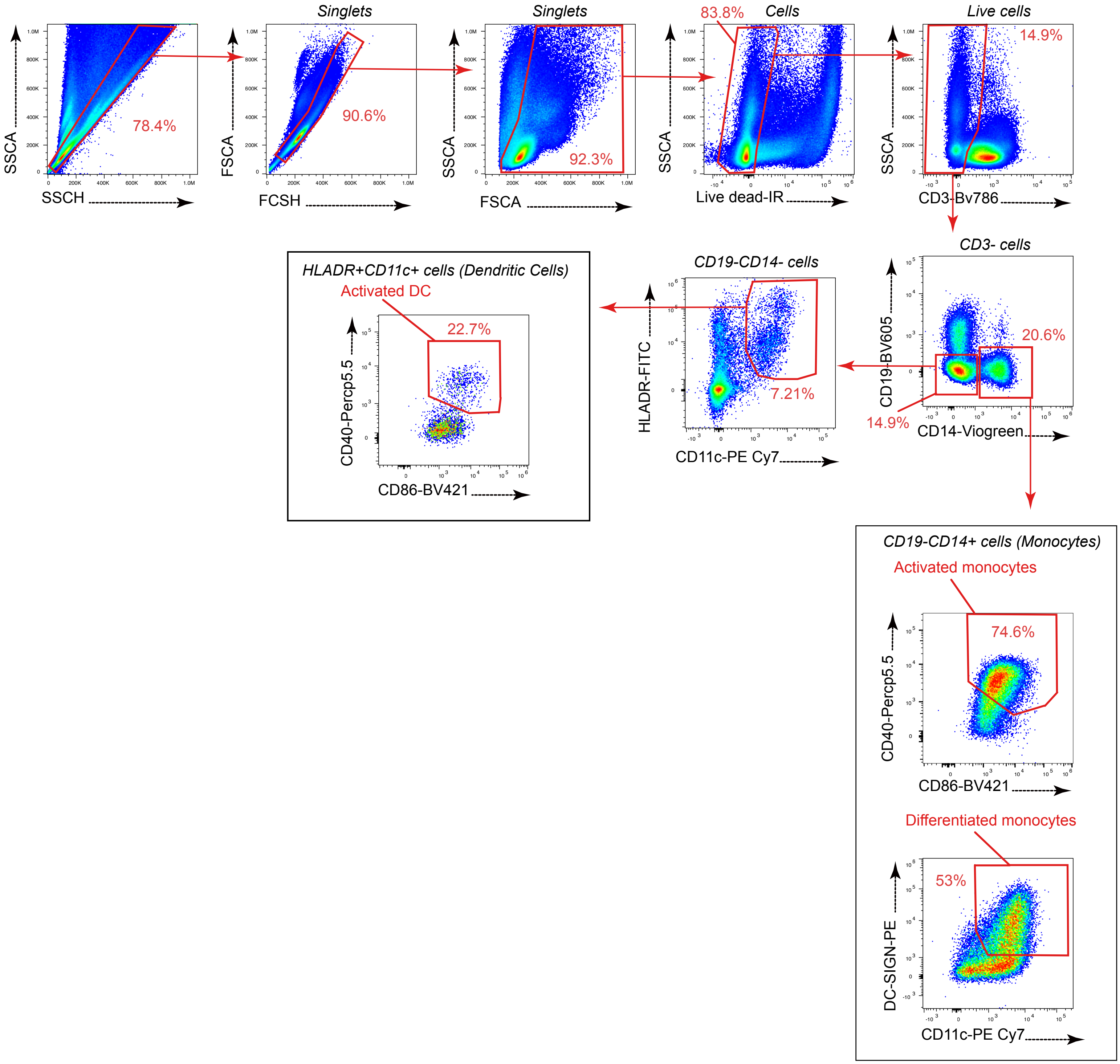


**Figure S5. Myeloid cell gating strategy.**
Monocytes were defined as CD3- CD19- CD14+ live singlet cells and were further phenotyped with the activation markers CD40 and CD86 or with the differentiation markers CD11c and DC-SIGN. Dendritic cells (DC) were defined as CD3- CD19- CD14- HLA-DR-hi CD11c-hi live singlets and were further phenotyped with the activation markers CD40 and CD86.

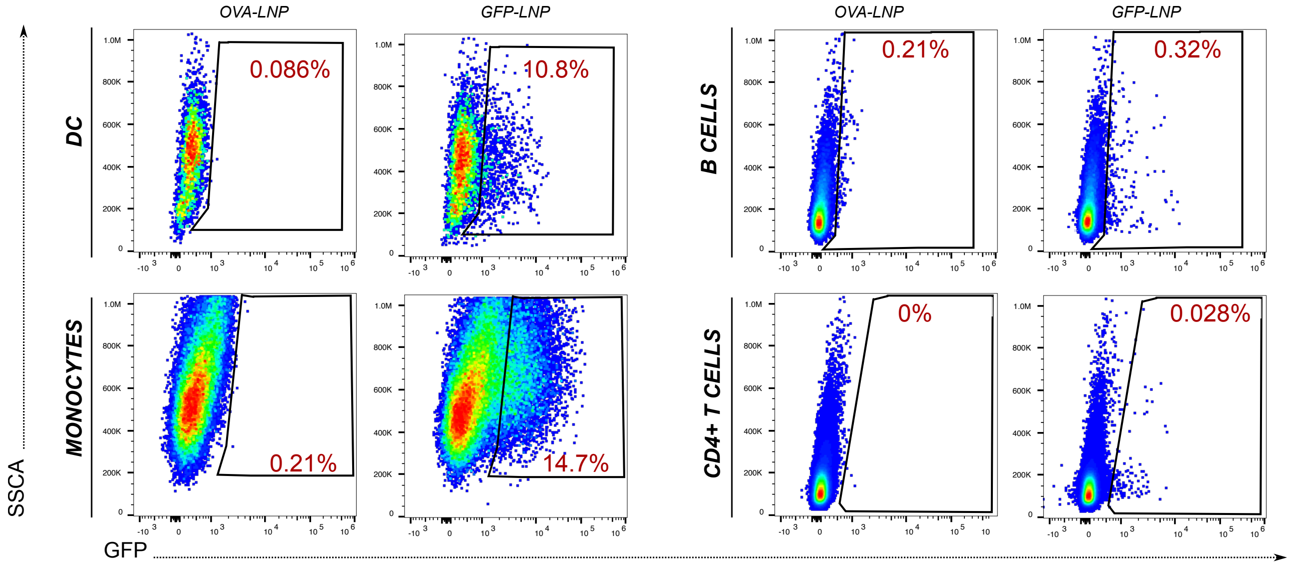


**Figure S6. Expression of a GFP-RNA lipid nanoparticle (LNP) in immune cell populations.** LO chips were perfused with a control LNP containing the ovalbumin mRNA (OVA-LNP) or with an LNP containing the reporter GFP mRNA (GFP-LNP). Cells were harvested from the LO chip at day 2 and the expression of GFP (on the x axis) was analyzed in the following cell populations: dendritic cells (DC, top left), monocytes (bottom left), B cells (top right), and CD4+ T cells (bottom right).

| Target | Fluorophore | Manufacturer | Reference | Dilution |
|---|---|---|---|---|
| **Flow cytometry** | | | | |
| Live dead | Aqua | Life technologies | L34966 | 800 |
| Live dead | IR | Life technologies | L34976 | 1000 |
| CD3 | Bv786 | BD bioscience | 565491 | 320 |
| CD4 | PE-CF594 | BD bioscience | 562281 | 200 |
| CD19 | Percp5.5 | BD bioscience | 561295 | 100 |
| CD19 | Bv605 | BD bioscience | 562653 | 100 |
| CD14 | v500 | BD bioscience | 561391 | 20 |
| CD11c | PE-cy7 | eBioscience | 25-0116-42 | 20 |
| HLADR | FITC | BD bioscience | 555811 | 20 |
| CD38 | APC-Cy7 | Biolegend | 353534 | 80 |
| CD27 | FITC | eBioscience | 11-0279-73 | 200 |
| IgG | Bv421 | Biolegend | 410704 | 320 |
| IgM | PE-Cy7 | Biolegend | 314532 | 320 |
| ICOS | AF647 | BD bioscience | 562834 | 100 |
| CD40 | Percp5.5 | Biolegend | 334316 | 20 |
| DC-SIGN | PE | Biolegend | 330106 | 20 |
| CD86 | v450 | BD bioscience | 560357 | 20 |
| IFN$\gamma$ | PE-Cy7 | Biolegend | 502528 | 100 |
| TNF$\alpha$ | Bv421 | Biolegend | 502932 | 100 |
| IL-2 | Percp5.5 | Biolegend | 500322 | 100 |
| Streptavidin | PE | Miltenyi | 130-106-789 | 20 |
| Streptavidin | AF647 | Invitrogen | S21374 | 20 |
| Ab102 (1 mg/ml) | AF647 | Homemade (C.P.) | / | 200 |
| mGO53 (1 mg/ml) | AF647 | Homemade (C.P.) | / | 200 |
| **Microscopy** | | | | |
| CD19 | AF561 | eBioscience™ | 505-0199-42 | 25 |
| CD4 | AF488 | BD biosciences | 557695 | 25 |
| ICOS | uncoupled | Cell Signaling Technology | 89601 | 100 |
| Ki67 | uncoupled | ThermoFisherScientific | MA514520 | 200 |
| Rabbit IgG | AF647 | Invitrogen | A32733 | 200 |

**Table S7: References of antibodies used in the study**
